# Mapping Individualized Dual-Axis Network Topology in Focal Epilepsy: Divergent Alterations in System Integrity, Integration, and Clinical Correlates

**DOI:** 10.64898/2026.03.17.712432

**Authors:** Qirui Zhang, Arielle Dascal, Sam S Javidi, Ankeeta Ankeeta, Michael R Sperling, Zhiqiang Zhang, Joseph I Tracy

## Abstract

Focal epilepsy is increasingly recognized as a disorder of distributed brain systems, yet characterizations of patient-specific alterations in network organization that are clinically meaningful remain limited. Here, we used resting-state functional MRI to introduce an overlap-permitting individualized network-estimation framework anchored to normative references to derive two complementary, system-level axes of topology: (1) network correspondence, indexing the expression of canonical networks and the fidelity of intrinsic system boundaries, and (2) *k*-hubness, indexing cross-system integration and multi-functionality through multi-network participation. We used a large presurgical focal epilepsy cohort (305 patients, including temporal and extratemporal epilepsy; 224 healthy participants) and an independent multi-syndrome validation cohort spanning focal and common generalized and childhood epilepsy syndromes (903 patients; 666 healthy participants), to show: (1) correspondence disruption of both canonical and non-normative networks constitutes a broadly shared system-level endpoint that displays heterogeneity across individuals and differential impacts on cognition, (2) *k*-hubness reconfigurations reflect lateralized and syndrome-dependent redistributions of cross-system integration, demonstrating epilepsy’s impact on the functional multiplicity inherent to specific brain areas. These complementary axes shared variance yet showed dissociable clinical associations, with correspondence preferentially relating to neurocognitive deficits and *k*-hubness aligning with epilepsy clinical features. Accordingly, these axes demonstrate the presence of distinct topological phenotypes, providing clinically relevant, reproducible patient-level signatures that explain both the syndrome-specific and common effects of seizures on the organization and integration of canonical and individualized brain systems.

## INTRODUCTION

Focal epilepsy is increasingly recognized as a disorder of distributed brain systems rather than an isolated focal pathology ^1, 2, 3, 4^. Yet clinical decision-making still hinges on patient-level questions: where the epileptogenic network resides, what brain systems the epilepsy has impacted, and who is at risk for cognitive morbidity or an unwanted treatment response. Despite advances in structural neuroimaging ^5, 6, 7^, these questions remain unanswered for a substantial subset of individuals. Resting-state functional MRI (rs-fMRI) and connectome studies have shown that epileptic activity perturbs large-scale functional architecture, extending beyond the presumed seizure focus to engage multiple interacting networks ^8, 9, 10, 11^. However, a central barrier to translating this knowledge into personalized clinical algorithms is the marked heterogeneity of these disturbances at the individual level. Although quantified at the group level, they vary substantially across patients and mirror the variability in seizure semiology, pathology, and neurocognitive deficits ^12, 13, 14^. Accordingly, the field is highly motivated to develop biomarkers of functional system architecture and potential network dysfunction that are applicable to individual patients and their planned treatments.

To characterize intrinsic network organization at the system level, clinically useful markers must quantify both **system integrity** (whether canonical functional systems and their spatial topography remain delineated as expected within an individual; whether highly individual systems exist with delineation of their link to disease) ^14, 15, 16^ and **system integration** (how brain regions participate in multiple systems and communicate through cross-network architecture, therefore serving multiple functionalities) ^17, 18, 19^(Fig. 1a). Conventional graph-theoretic metrics defined on fixed node–edge representations are constrained by the parcellation scheme imposed, enforcing single-network assignment^8, 9, 11, 20, 21, 22^, limiting sensitivity to the individual variation in system boundaries and ignoring the participation of multiple overlapping networks. To overcome these constraints, we utilized a robust sparse dictionary-learning decomposition that generated unique and reliable subject-specific network estimates, while accounting for voxel-wise overlap, indicative of multi-network participation. This method yielded a unified characterization of functional topology, capturing substrates of system-level pathology, going well beyond typical pairwise connectivity analyses^18, 19, 23, 24^.

**Figure 1.**
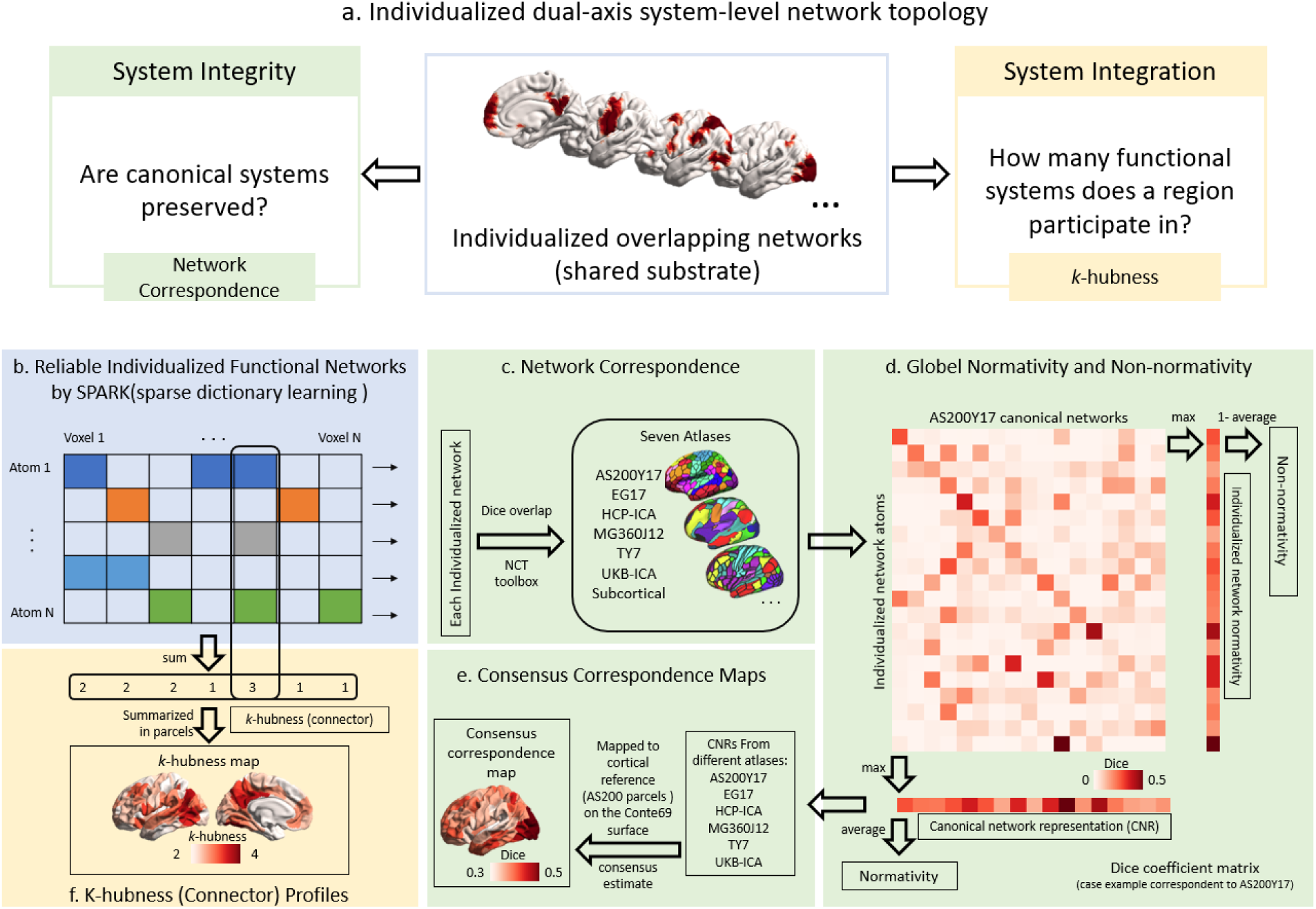
Conceptual framework and analytic pipeline. **(a)** Conceptual framework of individualized dual-axis system-level network topology. Individualized overlapping functional networks constitute a shared substrate from which two complementary but dissociable axes are derived: system integrity, indexing the preservation of canonical system boundaries, and system integration, indexing multi-network participation and cross-system connector topology. **(b)** Subject-specific resting-state functional networks (atoms) are estimated with a probabilistic dictionary-learning framework (SPARK). **(c)** Individualized networks are compared with canonical intrinsic connectivity network atlases. **(d)** Network correspondence is summarized using three complementary indices: normativity (overall alignment to canonical systems), non-normativity (idiosyncratic organization not explained by canonical architecture), and canonical network representation (CNR) (expression alignment of each canonical system within an individual). **(e)** To reduce dependence on any single template, correspondence measures are computed across multiple canonical atlases and aggregated into a cross-atlas consensus correspondence map capturing effects consistent across atlases. **(f)** *k*-hubness is computed from overlapping individualized networks to index multi-network participation and functionality, revealing connector-like topology beyond one-to-one template correspondence.

More specifically, we quantified two separate axes of brain system-level information. First, for system integrity, we quantified network correspondence by comparing every individual’s intrinsic networks against canonical intrinsic connectivity systems, and decomposed correspondence into normativity (alignment to canonical organization) and non-normativity (systematic, idiosyncratic organization not captured by canonical templates). Although rarely quantified explicitly, correspondence disruptions have been linked to cognitive integrity in temporal lobe epilepsy (TLE)^25, 26^, noting that such links to non-normative systems have come only from our prior work^25^. To reduce template-specific bias and spatially resolve and localize regions where correspondence is consistently disrupted, we constructed cross-atlas “consensus” correspondence maps. Second, to index system integration, we quantified *k*-hubness as a topology-based measure of multi-network participation and connector-like integration, capturing cross-system communication and the multi-functionality within regions going beyond one-to-one system correspondence. In this regard, we note that prior work in mesial-TLE indicates distinct syndromes based on seizure lateralization effects, which may relate in unique ways to the redistribution of integrative capacity^18, 19^.

An important unresolved question is the extent to which these two topological axes generalize beyond TLE to the broader focal epilepsy spectrum, including extratemporal epilepsy (EXE). Moreover, the translational value of these axes will hinge on whether they can be dissociated (i.e., differentially tracking neurocognitive functioning or providing linkages to clinically actionable seizure features such as lateralization and pathology), thereby providing specificity beyond simple estimates of disease severity or feature covariation. Equally important, clinical importance will depend upon demonstrated robustness across independent cohorts, across brain parcellation schemes, along with clarity on the specificity or generalized nature of the axes across common epilepsy syndromes^3, 5, 11, 27^.

Guided by the hypothesis that both canonical and individualized system-level network topology can be decomposed into complementary but distinct axes capturing functional-system integrity and cross-system integration, with each relating to distinct clinical features, we pursued three aims. First, we mapped normative topology and normative deviations in network correspondence and *k*-hubness in a large focal epilepsy cohort using MRI-based normative modeling, testing for syndrome-specific (TLE or EXE) and lateralization-specific (left or right) patterns. Second, we examined whether network correspondence and *k*-hubness exhibit dissociable multivariate associations with cognitive versus clinical features, and whether correspondence disruptions resolve into reproducible epilepsy subtypes and stages of severity that generate meaningful patient stratifications. Third, we replicated key focal-epilepsy findings in an independent cohort to assess cross-cohort stability and extend the analyses to other common epilepsy syndromes (genetic generalized epilepsy, GGE; self-limited epilepsy with centrotemporal spikes, SeLECTS; and absence epilepsy). This allowed us to evaluate for specificity to focal epilepsy and produce spectrum level depictions of topology and brain system organization across epilepsy syndromes.

## RESULTS

### Study design

We analyzed two rs-fMRI cohorts to quantify individualized functional network topology and test for both syndrome-specificity and cross-cohort generalizability. Our primary discovery cohort comprised a presurgical focal epilepsy sample (305 patients, age range: 15-73 years, including 249 with TLE and 56 with EXE) from Thomas Jefferson University (TJU dataset), along with a healthy participant (HP) comparison group (224 HPs, age range: 18-65 years). The independent validation cohort from Jinling Hospital (JLH dataset; 903 patients, age range: 6-56 years, including 659 with focal epilepsy, 108 with GGE, 112 with SeLECTS, and 24 with absence epilepsy; plus 666 matched HPs, age range: 4-70 years) was intentionally broader in syndrome composition. This design allowed us to test whether focal epilepsy–related findings replicate at an independent site under different clinical ascertainment and imaging conditions, and whether the same analytic framework and findings generalized to other common epilepsy syndromes. Sample demographic and clinical characteristics are summarized in Supplementary Tables 1 (TJU) and 2 (JLH).

To characterize individualized network topology, we quantified two distinct but complementary classes of imaging markers: **network correspondence**, capturing how an individual’s data-driven networks relate to established, canonical intrinsic connectivity systems, and ***k*-hubness**, capturing connector-like properties via multi-network participation within individualized networks. For network correspondence, we used SParsity-based Analysis of Reliable *k-*hubness (SPARK), a probabilistic dictionary-learning framework to derive a unique data-driven functional network set for each individual (Fig. 1b), which we then compared to the canonical intrinsic connectivity networks (Fig. 1c), and calculated the following four complementary summary measures (Fig. 1d). (i) **Normativity** indexed overall alignment to canonical systems, (ii) **non-normativity** quantified idiosyncratic organization not explained by canonical architectures, and (iii) **canonical network representation (CNR)** captured the degree to which each canonical system was expressed in a given individual. To minimize dependence on any single canonical atlas, these correspondence measures were computed across multiple widely used, openly available network atlases, allowing us to construct a (iv) **cross-atlas consensus correspondence map** to verify if effects were consistent across atlases. This reduced template-specific bias and highlighted robust network alterations that generalize across different spatial resolutions of brain architecture (Fig. 1e). We then quantified ***k*-hubness** from overlapping individualized networks to index multi-network participation and functionality, capturing connector-like topology and inter-network integration/communication beyond one-to-one correspondence with canonical systems (Fig. 1f). To enable fair comparison across cohorts and to reduce confounding from demographic and acquisition-related factors, all network- and region-level metrics were converted to **covariate-adjusted normative deviation scores (*w*-scores)** by estimating expected values from each cohort’s healthy participants and expressing each individual’s residual in units of healthy variability.

### Network correspondence of individualized brain network topology

Across canonical atlases, individuals with focal epilepsy exhibited a marked reduction in mean **normativity**, indicating diminished alignment between individualized networks and canonical intrinsic connectivity systems (mean normativity = −0.356 ± 1.200, *P* < 0.001, FDR-corrected; Fig. 2a). This reduction was consistently observed across hierarchical focal epilepsy subgroups, including the overall focal epilepsy cohort, the temporal lobe epilepsy (TLE) and extratemporal epilepsy (EXE) subgroups within it, and the left- and right-sided TLE subgroups defined within the TLE group. In parallel, focal epilepsy was characterized by a significant increase in mean **non-normativity** (mean non-normativity = 0.404 ± 1.072, *P* < 0.001, FDR-corrected; Fig. 2b), again consistently present across all subgroups, reflecting enhanced individualized or redistributed network organization beyond canonical architectures. Atlas-specific analyses are provided in Supplementary Fig. 1. While the overall pattern was robust across atlases, their discriminative sensitivity differed: the EG17 template provided the strongest separation for normativity, whereas the UKB-ICA template was most sensitive to non-normativity, yielding the largest deviations relative to HPs. In contrast, the Yeo 7-network atlas showed weaker patient discrimination from HPs, likely due to its relatively coarse spatial granularity.

**Figure 2.**
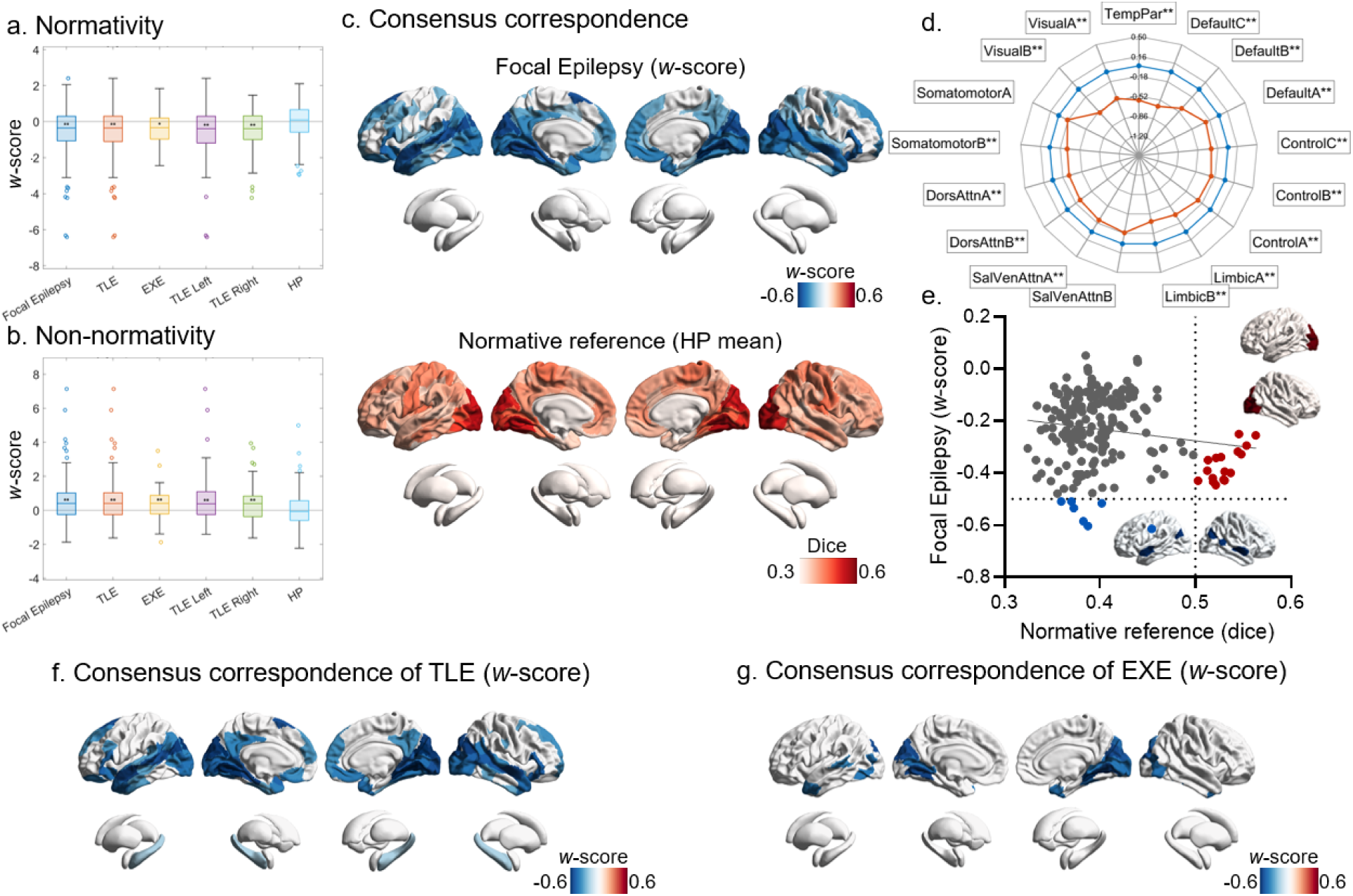
Network correspondence disruptions in focal epilepsy. Group-level deviations in mean normativity (a) and mean non-normativity (b), expressed as *w*-scores and aggregated across canonical atlases (\**P* < 0.05, \*\**P* < 0.01, FDR corrected).

Spatially resolved consensus correspondence maps were obtained by aggregating atlas-specific results across cortical parcels defined in the AS200^28^ atlas and evaluated using permutation-based max-T correction across regions or networks (*P* < 0.05), revealed widespread correspondence reductions in focal epilepsy. Regionally, decreases were most prominent in lateral temporal, occipital/visual, and frontal cortices (Fig. 2c). When summarized across the 17 large-scale networks, significant reductions were observed in all networks except Somatomotor A and Salience/Ventral Attention B, with the largest effects in Default C, Temporal Parietal, and Visual A and B networks (Fig. 2d). The normative reference brain map, showing the mean raw consensus correspondence in HPs, indicated higher correspondence in unimodal regions, particularly visual and somatomotor cortex (Fig. 2c). A spatial comparison between the focal epilepsy deviation map and the normative reference indicated a weak inverse association (*r* = −0.16, *P* = 0.022, uncorrected; Fig. 2e). Notably, the most extreme reductions clustered in bilateral temporal cortex. In addition, regions with high normative correspondence in HP, especially visual cortex, also exhibited substantial reductions, indicating that correspondence disruption is not restricted to regions with low baseline correspondence.

Subgroup analyses demonstrated both shared and syndrome-specific patterns. TLE showed a reduction profile that broadly mirrored the overall focal epilepsy pattern, with additional bilateral weak hippocampal involvement (Fig. 2f). In contrast, EXE exhibited a more restricted pattern, with reductions largely confined to core networks including Default C, Temporal Parietal, and Visual A and B networks (Fig. 2g). Atlas specific analyses are provided in Supplementary Fig. 2.

Normativity summarizes overall alignment between individualized networks and canonical intrinsic connectivity systems, whereas non-normativity captures individualized network organization not explained by canonical architectures. (c) Spatial consensus correspondence maps for focal epilepsy (*w*-scores), derived by aggregating atlas-specific results, highlighting cortical regions showing significantly reduced correspondence (blue) in focal epilepsy and in each focal-epilepsy subgroup (*P* < 0.05). The normative reference brain map displays the mean raw consensus correspondence in healthy participants (HP). (d) The accompanying radar plot summarizes correspondence effects across the 17 large-scale networks. The orange line denotes focal epilepsy, and the blue line marks the *w*-score = 0 reference line (\**P* < 0.05, \*\**P* < 0.01). (e) The scatter plot shows that the consensus correspondence deviation map in focal epilepsy exhibits only a weak spatial association with the normative reference map (*r* = −0.16, *P* = 0.022, uncorrected). Notably, regions with the most pronounced reductions in correspondence are localized to the bilateral temporal cortex (blue points and blue-highlighted regions). In addition, the visual cortex, which exhibits among the highest correspondence in the normative reference (red points and red-highlighted regions), also showed substantial correspondence reductions, indicating that even systems with high baseline correspondence are affected. Syndrome-specific analyses show consensus correspondence maps for TLE (f) and EXE (g) in *w*-scores (*P* < 0.05). Group-level normativity and non-normativity statistics are corrected using FDR, whereas consensus correspondence results are assessed using permutation-based max-T correction.

### Subtype and Staging of Network Correspondence Disruptions

Because patients with focal epilepsy showed a largely unidirectional disruption in network correspondence, we used the Subtype and Stage Inference (SuStaIn) framework to decompose this disruption into putative subtypes and stages of evolving decline and severity. Two correspondence-disruption subtypes were identified (Subtype 1: *n* = 104; Subtype 2: *n* = 79). Individuals assigned to Stage 0 were classified as the “network correspondence normal” group (*n* = 115), indicating that none of their regional correspondence *z*-scores fell below −1 (i.e., no progression). Subtypes 1 and 2 exhibited distinct stage-dependent progression patterns. In Subtype 1, early involvement was centered on default-mode regions with a temporal-lobe predominance, followed by expansion toward frontoparietal systems and ultimately widespread involvement including Somatomotor and Visual cortices. In contrast, Subtype 2 preferentially involved visual cortex and Somatomotor networks at early stages, then propagated toward frontoparietal systems, and eventually became widespread, including default-mode regions (Fig. 3a). When the 17 canonical networks were ordered along the principal functional connectivity gradient^29, 30^, this dissociation was more clearly expressed: Subtype 1 was biased toward the transmodal end of the gradient, whereas Subtype 2 was biased toward a unimodal origin; each subtype, however, clearly converged on a common endpoint involving a pan-network breakdown in correspondence along the gradient axis (Fig. 3b).

**Figure 3.**
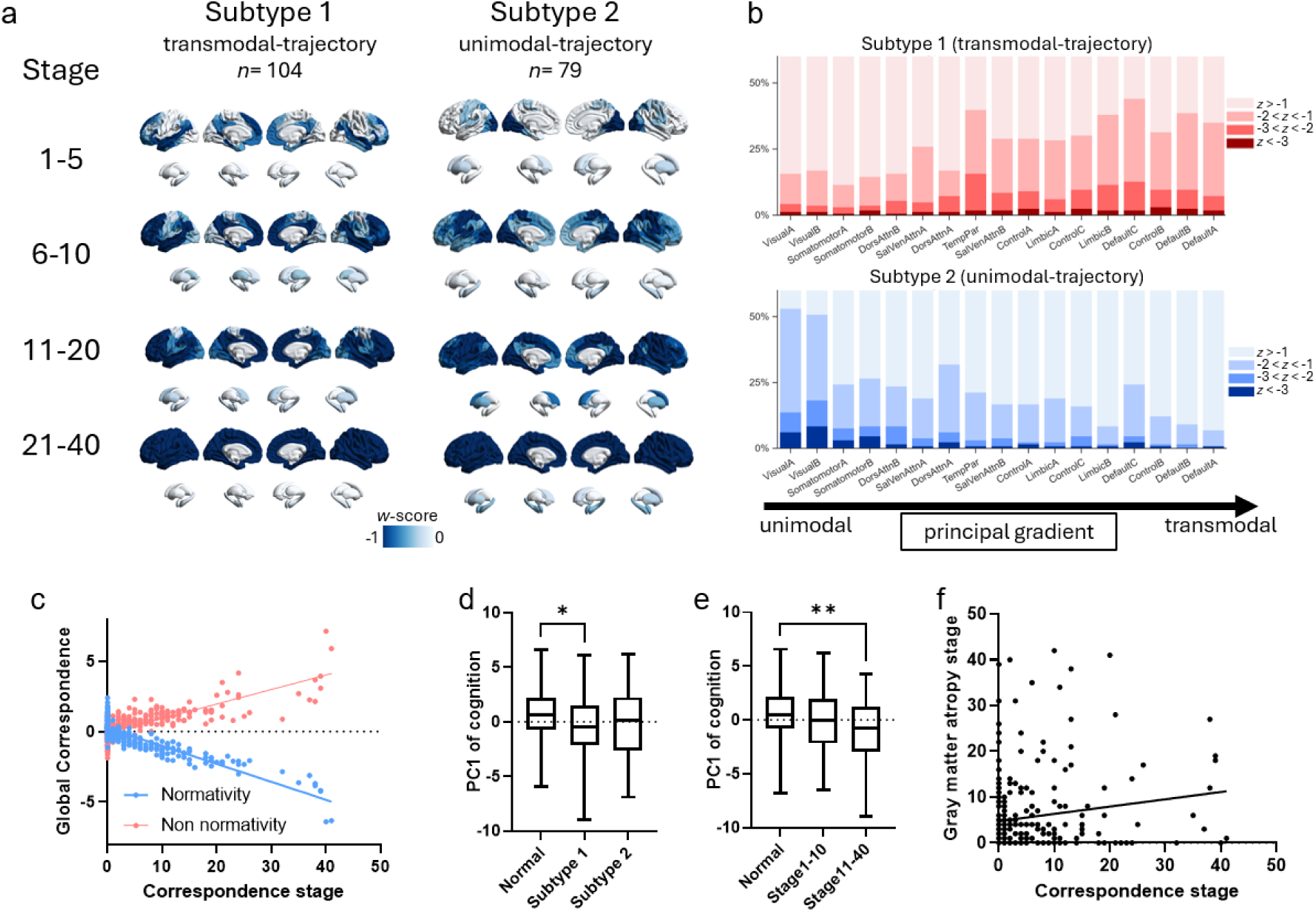
Subtypes and staging of network correspondence disruptions in focal epilepsy. (a) Mean regional *w*-scores of network correspondence disruptions across inferred stages for the two SuStaIn-derived subtypes. (b) Stacked bar plots showing the proportion of correspondence disruptions within each of the 17 canonical networks for the two subtypes, ordered along the principal functional connectivity gradient. Deviations are displayed using three thresholds (*z*[*w*-score] = 1, 2, and 3). Subtype 1 is biased toward pan-network/transmodal systems (transmodal-trajectory), whereas Subtype 2 is biased toward unimodal systems (unimodal-trajectory). (c) Scatter plots illustrating associations between correspondence disruption stage and global correspondence metrics, showing a strong negative correlation with normativity and a strong positive correlation with non-normativity. (d) Box plots comparing cognitive performance summarized by the first principal component (PC1) across the network correspondence–normal group, Subtype 1, and Subtype 2, demonstrating significantly reduced cognition at subtype 1 (* *P* < 0.05, Tukey-corrected). (e) PC1 is also compared across the network correspondence–normal group, early stages (Stage 1–10), and late stages (Stage 11–40), demonstrating significantly reduced cognition at later stages (** *P* < 0.01, Tukey-corrected). (f) Relationship between network correspondence disruption stage and gray matter volume atrophy stage derived from a parallel atrophy-based SuStaIn analysis.

The stage pattern revealed the expected associations with global correspondence metrics: stage correlated strongly and negatively with normativity (*r* = −0.89, *P* < 0.001) and positively with non-normativity (*r* = 0.80, *P* < 0.001) (Fig. 3c). Cognition was summarized using the first principal component (PC1) across all neuropsychological measures, which explained 35.5% of the variance. Compared with the correspondence-normal group, Subtype 1 showed significantly lower cognition, whereas Subtype 2 did not differ significantly (one-way ANOVA: *F* = 3.56, *P* = 0.029; Tukey-corrected: Normal vs subtype 1, mean difference = 1.03, *P* = 0.015; Normal vs subtype 2, mean difference = 0.51, *P* = 0.413) (Fig. 3d). When patients were instead stratified by correspondence-disruption stage irrespective of subtype, cognition showed a clearer stage dependence (one-way ANOVA: *F* = 5.107, *P* = 0.006), driven by significantly lower PC1 in late stages (Stages 11–40) relative to the correspondence-normal group (Tukey-corrected: mean difference = 1.553, *P* = 0.0045), whereas early stages (Stages 1–10) did not differ (mean difference = 0.433, *P* = 0.468) (Fig. 3e). In contrast, correspondence subtypes showed no significant differences across clinical characterization variables, including epilepsy type, focal-to-bilateral tonic–clonic seizures (FBTCS), seizure lateralization, or underlying pathology (Supplementary Fig. 3c).

While functional staging of network organization has not previously been reported, structural staging, most commonly based on gray matter decline, has been examined in a prior study^12^. We therefore sought to determine whether these two forms of staging are related. To accomplish this, in parallel, we performed a gray matter volume-based SuStaIn analysis following prior work and recovered three atrophy patterns consistent with the literature ^12^. Notably, the stage of network correspondence disruption was not associated with gray matter volume (GMV) atrophy stage (Spearman *r* = 0.07, *P* = 0.22, Fig. 3f), and collinearity between the two staging estimates was minimal (VIF = 1.025), indicating negligible shared variance. The gray matter volume SuStaIn results are shown in Supplementary Fig. 3a-b.

### *k*-hubness of individualized brain network topology

*k*-hubness maps were obtained by aggregating voxel-level maps into AS200^28^ parcels and subcortical regions. Patients with focal epilepsy showed marked alterations in *k*-hubness, indicating disrupted multi-network participation and connector-like topology. Relative to healthy participants (HP), *k*-hubness was significantly reduced in the bilateral temporal poles, superior temporal gyri, and hippocampi (Fig. 4a), consistent with diminished multi-network integration capacity within temporo-limbic circuitry. In contrast, *k*-hubness increases were observed in the bilateral visual cortices and the right dorsolateral prefrontal cortex, suggesting a redistribution of integrative load toward sensory and association regions. Figure 4a also includes a normative reference map of mean raw *k*-hubness in HP, which showed the expected pattern of higher *k*-hubness in transmodal cortex and lower *k*-hubness in unimodal systems.

**Figure 4.**
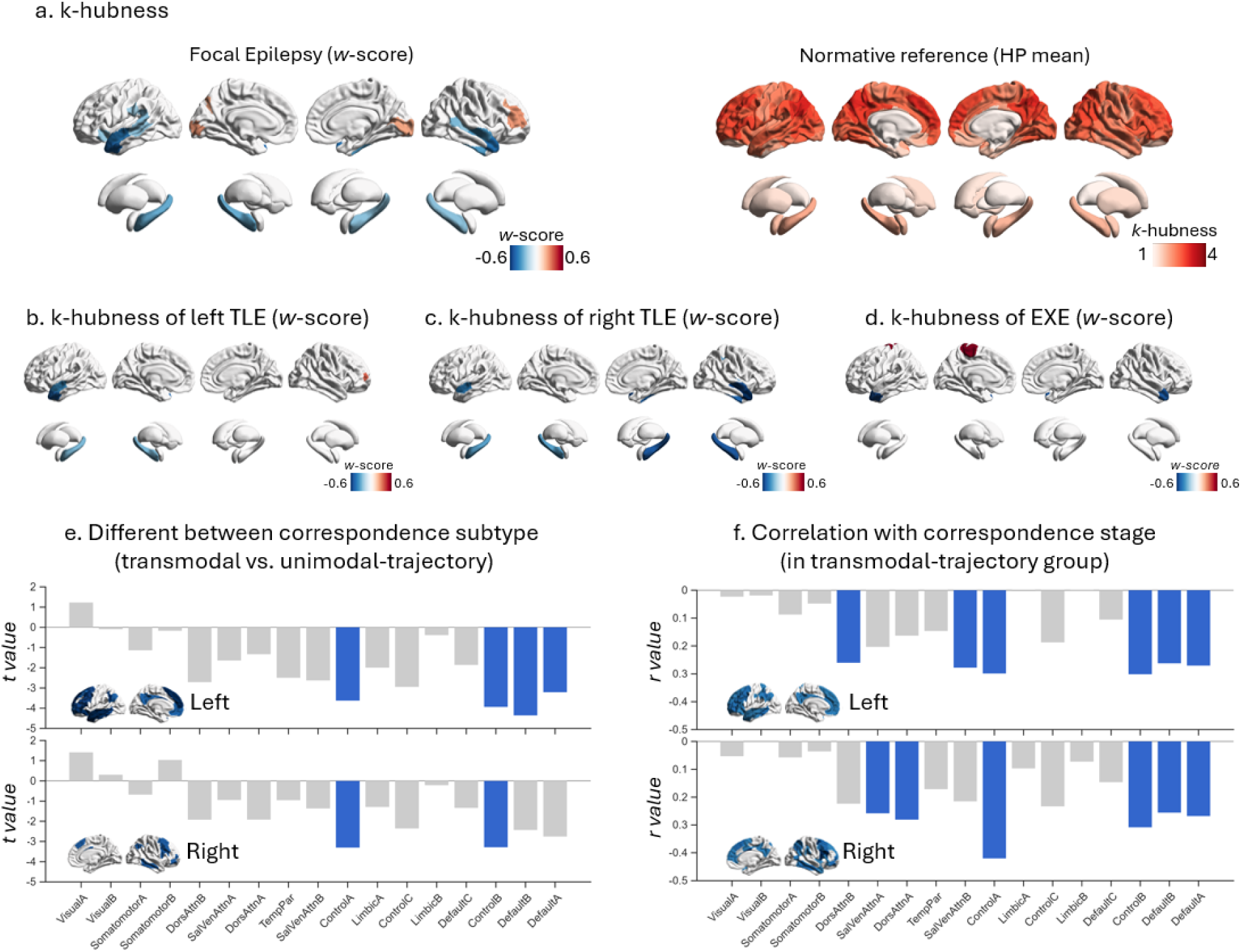
Altered *k*-hubness reveals disrupted multi-network integration in focal epilepsy. (a) Spatial maps of *k*-hubness alterations in focal epilepsy relative to HP, expressed as *w*-scores (left). Focal epilepsy is characterized by significant reductions in *k*-hubness in the bilateral temporal poles, superior temporal gyri, and bilateral hippocampi. The normative reference brain map displays the mean raw *k*-hubness in HP (right). (b) Left TLE and (c) right TLE show pronounced lateralization and syndrome specificity of *k*-hubness alterations, predominantly manifesting as reductions in the ipsilateral temporal lobe and hippocampus. (d) EXE is characterized by reduced *k*-hubness in the bilateral temporal poles and increased *k*-hubness in the left motor cortex. (e) Bar plots show network-level differences in *k*-hubness (*t* values) between correspondence subtypes; blue bars indicate significant networks, corresponding to the blue regions in the brain maps, and are predominantly transmodal. (f) Within the transmodal-trajectory group, bar plots show network-level correlations between correspondence stage and *k*-hubness; blue bars indicate significant networks (matching the blue regions in the brain maps), again concentrated in transmodal systems. All displayed regions/networks survive permutation-based max-T correction (*P* < 0.05).

Subgroup analyses further demonstrated pronounced lateralization and syndrome specificity. Left TLE was characterized by *k*-hubness reductions predominantly in the ipsilateral temporal lobe and hippocampus (Fig. 4b), whereas right TLE showed a similar pattern with clear rightward predominance (Fig. 4c). EXE exhibited reduced *k*-hubness in the bilateral temporal poles alongside increased *k*-hubness in the left motor cortex (Fig. 4d), consistent with syndrome-dependent reconfiguration of multi-network integration.

At the network level, *k*-hubness differences between correspondence subtypes were concentrated in transmodal systems (Fig. 4e), indicating that patients classified as correspondence transmodal-predominant (trajectory) also showed preferential *k*-hubness alterations within transmodal networks. Moreover, within the transmodal-trajectory subgroup, correspondence stage was significantly associated with network-level *k*-hubness across a similar set of transmodal networks (Fig. 4f). Together, these findings suggested that correspondence-based phenotyping captures a broader transmodal axis of network reorganization that is mirrored by changes in multi-network participation, potentially reflecting both loss of integrative capacity in temporo-limbic hubs and compensatory or redistributed connector load across remaining large-scale systems.

### Associations with clinical-cognitive features

Sparse canonical correlation analysis (sCCA) identified multivariate relationships linking deviations in individualized network topology (captured by both correspondence metrics and *k*-hubness metrics, set X) to clinical and cognitive expressions (set Y). Using 1000 permutations, we detected a significant overall association between the two sets (*P* = 0.014). The first canonical variate (*r* = 0.33) primarily represented a correspondence profile characterized by increased non-normativity, reduced overall normativity, and reduced normativity within several large-scale systems (e.g., somatomotor, salience, and ventral-attention systems). This correspondence-dominant variate was coupled most strongly with better cognitive performance, including measures of verbal learning (e.g., California Verbal Learning Test), processing speed (Coding), and IQ (Fig. 5a). The second canonical variate (*r* = 0.37) was dominated by *k*-hubness-related features, including increased *k*-hubness in the right caudate and putamen, and reduced *k*-hubness across several large-scale networks (e.g., the control network). This variate aligned most closely with epilepsy-related clinical characteristics, including seizure lateralization, unilateral or bilateral hippocampal sclerosis (Fig. 5b).

**Figure 5.**
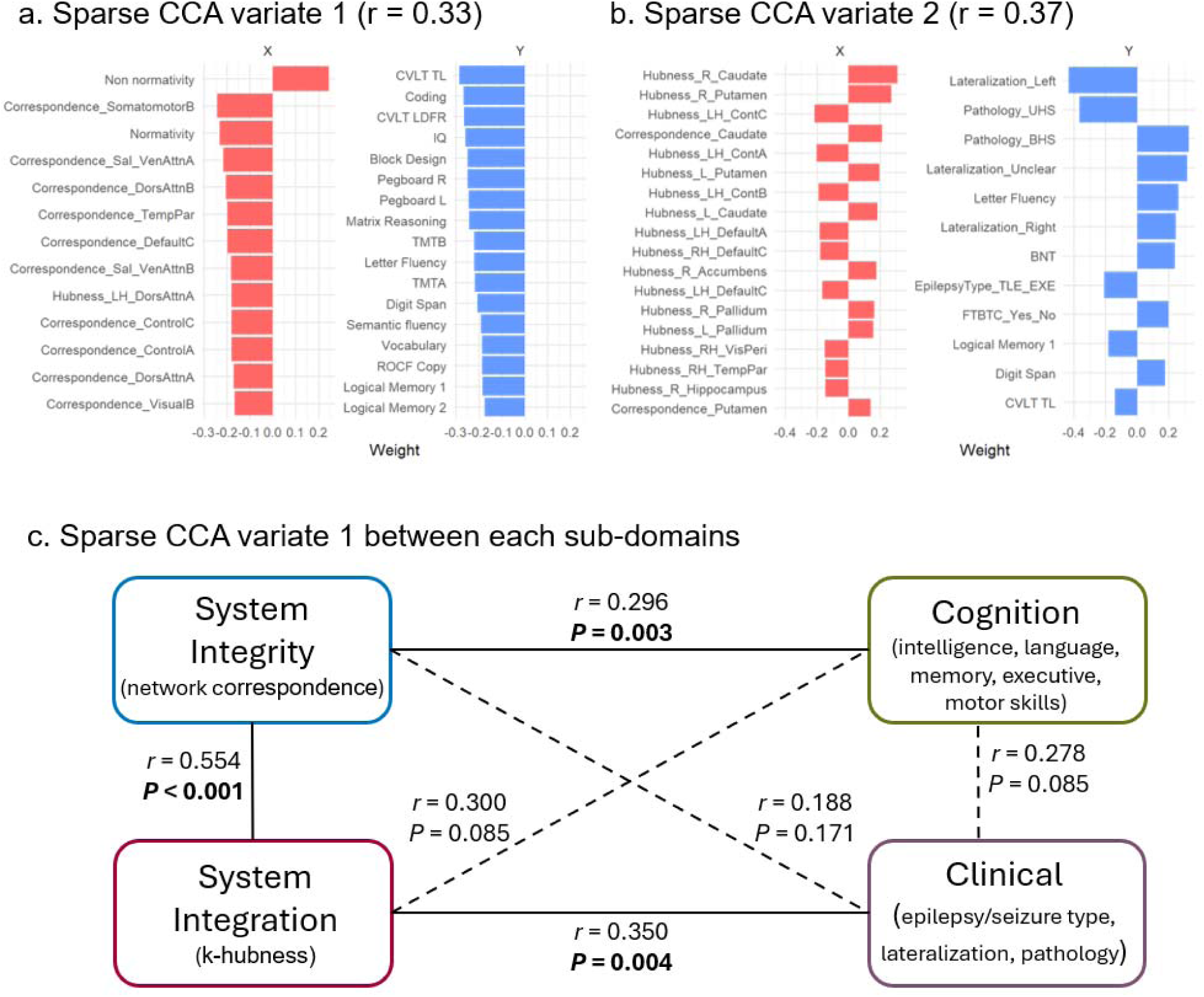
Sparse canonical correlation analysis (sCCA) between network topology and clinical–cognitive phenotypes. **(a)** First canonical variate (Variate 1) and **(b)** second canonical variate (Variate 2) from sCCA, showing the relative weights of imaging-derived network topology features (X set) and clinical–cognitive variables (Y set). Bar plots indicate the contribution of each feature to the variate. **(c)** Subdomain-level sCCA. The full feature space is partitioned into four a priori subdomains: System Integrity (network correspondence metrics), System Integration (*k*-hubness metrics), Cognition, and Clinical. Pairwise sCCA is performed across all subdomain pairs, and significance is evaluated using permutation-derived P values followed by BH-FDR correction across all tested domain pairs. Solid lines indicate significant associations after correction; dashed lines indicate non-significant associations.

To further resolve these dissociations, we conducted subdomain-level sCCA to isolate domain-specific relationships. We partitioned the full feature sets into four a priori subdomains: System Integrity, indexed by network correspondence metrics; System Integration, indexed by *k*-hubness metrics; Cognition, comprising intelligence, language, memory, executive, and motor domains; and Clinical, comprising epilepsy/seizure type, lateralization, and pathology (Fig. 5c). Pairwise sCCA was then performed across all subdomain pairs, and significance was evaluated using permutation-derived P values followed by BH-FDR correction across all tested subdomain pairings. System Integrity was significantly associated with Cognition (*r* = 0.296, *P* = 0.003) but not with Clinical features (*r* = 0.188, *P* = 0.171). Conversely, System Integration was significantly associated with Clinical features (*r* = 0.350, *P* = 0.004) but not with Cognition (*r* = 0.300, *P* = 0.085). System Integrity and System Integration were also significantly coupled (*r* = 0.554, *P* < 0.001), indicating shared variance across these complementary topology axes. In contrast, Cognition and Clinical subdomains were not significantly associated at the global level (*r* = 0.278, *P* = 0.085). Together, these subdomain-level results reveal correspondence metrics primarily reflect system-level cognitive consequences of epilepsy, whereas *k*-hubness more specifically indexes cross-system integration via multi-network participation. It should be noted that not all aspects of clinical severity or cognitive functioning were available or modeled in the present analysis; therefore, these multivariate associations are likely to capture only a subset of the full expressed clinical phenotype. We additionally performed more fine-grained, exploratory subdomain analyses to clarify which higher-resolution domains drove these relationships (Supplementary Fig. 4).

### Independent Cohort Validation

The main findings derived from the TJU cohort were robustly replicated in a completely independent dataset from another center (the JLH cohort). Group-level statistical maps for both network correspondence and *k*-hubness showed highly consistent patterns in the JLH dataset, including decreased normativity and increased non-normativity in focal epilepsy and its subgroups (Supplementary Fig. 5).

Consensus correspondence maps further revealed that focal epilepsy and its subtypes were characterized by the most pronounced reductions in correspondence within temporal, occipital, and frontal cortices (Fig. 6a). Spatial similarity, quantified by Pearson correlation between the statistical maps derived from the two datasets, was high for focal epilepsy (*r* = 0.76), TLE (*r* = 0.73), and EXE (*r* = 0.75), with all spin-test–corrected P values < 0.001 (Fig. 6a).

**Figure 6.**
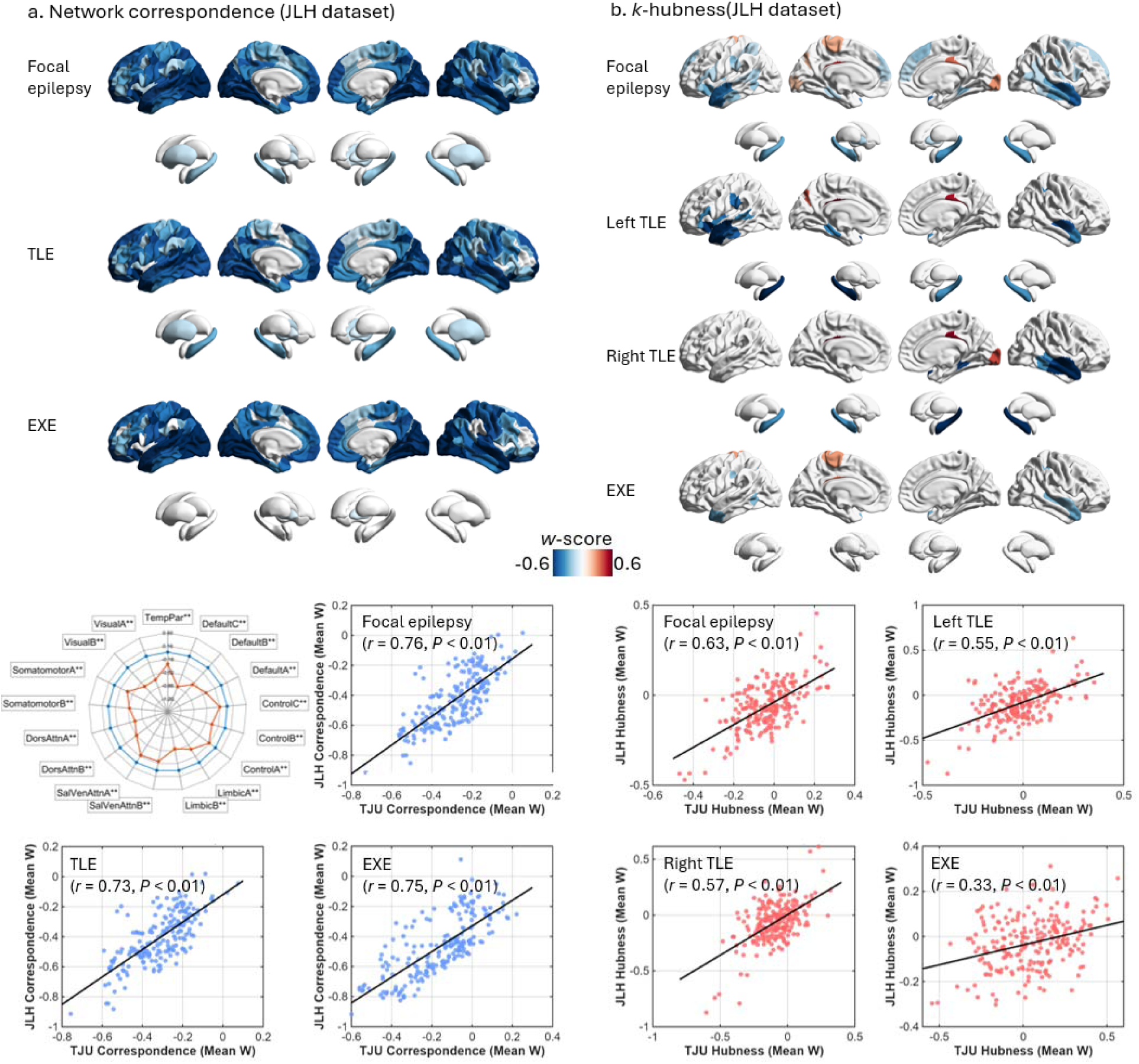
Independent cohort validation. (a) Network correspondence replication in an independent cohort (JLH). Brain maps show mean network correspondence *w*-scores for focal epilepsy and its subtypes, with a radar plot summarizing effects across the 17 large-scale networks for focal epilepsy (\*\**P* < 0.01). The scatter plot depicts the spatial similarity between TJU- and JLH-derived correspondence maps (Pearson correlation; spin-test-corrected). (b) *k*-hubness replication in the JLH cohort. Brain maps show mean *k*-hubness *w*-scores for focal epilepsy and its subtypes, with the scatter plot depicting spatial similarity between TJU- and JLH-derived *k*-hubness maps (Pearson correlation; spin-test-corrected). All brain maps display only regions surviving permutation-based max-T correction (*P* < 0.05).

Convergent replication was also observed for *k*-hubness alterations. In focal epilepsy, *k*-hubness was significantly reduced in bilateral temporal cortices and hippocampi. Pronounced lateralized reductions were evident in left- and right-sided TLE, whereas EXE was distinguished by increased *k*-hubness in the left motor cortex (Fig. 6b). These *k*-hubness patterns exhibited strong spatial similarity across datasets (focal epilepsy: *r* = 0.63; left TLE: *r* = 0.55; right TLE: *r* = 0.57; EXE: *r* = 0.33; all spin-test–corrected P values < 0.0001; Fig. 6b).

### The Distinct Network Topology of Focal Epilepsy and Other Epilepsy Syndromes

Leveraging the unique inclusion criteria of the JLH dataset, we extended our network topology framework to other common epilepsy syndromes, including predominantly adolescent and adult generalized genetic epilepsy (GGE), as well as pediatric epilepsies such as self-limited epilepsy with centrotemporal spikes (SeLECTS) and absence epilepsy (AE). This broader comparative analysis enabled us to determine whether the network-topological alterations observed in focal epilepsy reflect syndrome-specific features or shared properties across the epilepsy spectrum. At the global level, both GGE and SeLECTS showed a marked reduction in mean normativity and an increase in mean non-normativity, whereas AE showed no significant global change (Supplementary Fig. 5). At the regional level, both GGE and SeLECTS exhibited spatially scattered reductions in network correspondence; however, these changes lacked the clear regional specificity and coherent spatial organization observed in focal epilepsy. In contrast, AE showed abnormally increased network correspondence in bilateral thalamus and bilateral primary motor cortices, while SeLECTS demonstrated increased correspondence predominantly in the bilateral thalamus. These findings likely reflect enhanced cortico–thalamic coupling associated with generalized or developmental epileptic activity patterns in AE and SeLECTS (Fig. 7a). With respect to *k*-hubness, only SeLECTS displayed significant reductions, primarily involving prefrontal regions (Fig. 7b), whereas other syndromes showed no robust *k*-hubness alterations comparable to focal epilepsy.

**Figure 7.**
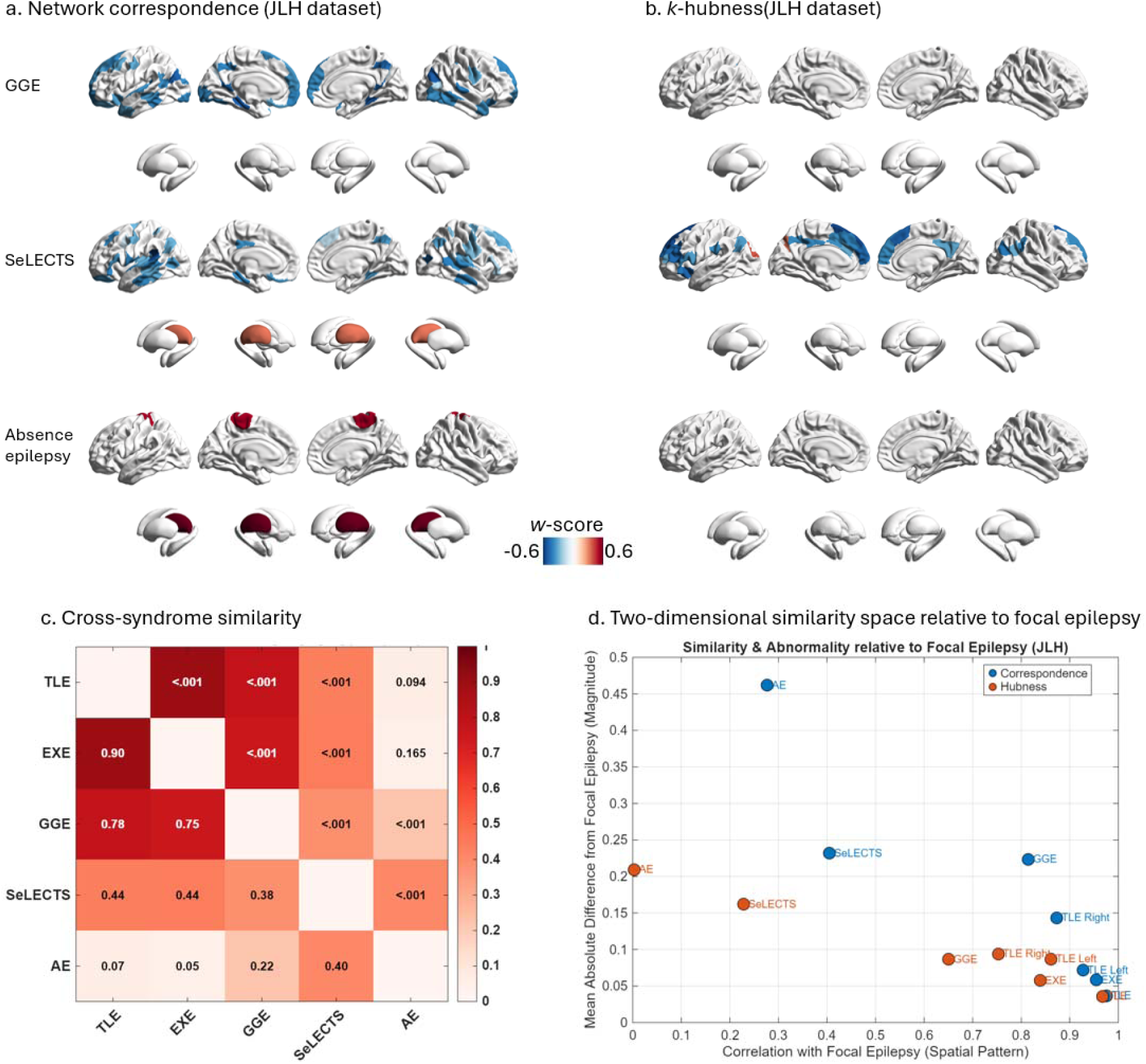
Distinct network topology of focal epilepsy across epilepsy syndromes. (a) Network correspondence *w*-score maps in the JLH cohort for generalized genetic epilepsy (GGE), self-limited epilepsy with centrotemporal spikes (SeLECTS), and absence epilepsy. (b) *k*-hubness *w*-score maps for the same syndromes. All brain maps display only regions surviving permutation-based max-T correction (*P* < 0.05). (c) Cross-syndrome similarity heatmap based on group-mean topology features (combined correspondence and *k*-hubness). The lower triangle shows Pearson correlation coefficients, and the upper triangle shows spin-test–corrected P values. (d) Two-dimensional similarity space relative to focal epilepsy, with the x-axis representing spatial pattern similarity (correlation with focal epilepsy) and the y-axis representing magnitude deviation (mean absolute difference). Red points indicate *k*-hubness and blue points indicate network correspondence; positions closer to the lower-right corner denote greater similarity to focal epilepsy.

To quantify cross-syndrome similarity in network topology, we integrated correspondence and *k*-hubness metrics into a unified feature space comprising 421 topology-derived features. Group-mean feature vectors were then compared across syndromes using Pearson correlation, with spatial autocorrelation accounted for using spin tests. TLE and EXE exhibited extremely high similarity (*r* = 0.90, *P* < 0.001), confirming their close topological relationship within focal epilepsy. GGE also showed strong similarity to both TLE (*r* = 0.78, *P* < 0.001) and EXE (*r* = 0.75, *P* < 0.001), suggesting partial convergence in large-scale network organization despite distinct clinical phenotypes. In contrast, SeLECTS demonstrated only moderate similarity to adult epilepsies and AE (TLE: *r* = 0.44; EXE: *r* = 0.44; GGE: *r* = 0.38; AE: *r* = 0.40; all *P* < 0.01), while AE and GGE—both classified as generalized epilepsies—showed only weak similarity (*r* = 0.22, *P* < 0.01) (Fig. 7c).

To further contextualize these relationships, we projected each syndrome into a two-dimensional similarity space relative to focal epilepsy, where the x-axis represented spatial pattern similarity (correlation with focal epilepsy) and the y-axis represented magnitude deviation (mean absolute difference from focal epilepsy). In this space, TLE and EXE clustered closest to focal epilepsy, occupying the lower-right region characterized by high spatial similarity and low magnitude difference. GGE occupied an intermediate position, followed by SeLECTS, which showed moderate similarity but larger deviations. AE was positioned furthest from focal epilepsy, indicating both low spatial similarity and large topological divergence (Fig. 7d). Collectively, these results demonstrated that focal epilepsy is characterized by a distinctive and coherent network-topological profile that is not shared uniformly across other epilepsy syndromes. While partial overlap exists, particularly with GGE, the combination of correspondence and *k*-hubness metrics revealed a graded separation across syndromes, underscoring the specificity of focal epilepsy within the broader epilepsy spectrum.

### Sensitivity analyses

Several sensitivity analyses were conducted to assess the robustness of the main findings. First, because individualized networks derived from SPARK may contain noise-related atoms^18, 19^, we performed an explicit noise-identification procedure. Each atom was visually inspected and classified as signal or noise using previously described criteria (e.g., atom located along brain boundaries or outside the brain, as well as vascular- and motion-related artifacts) ^31^. In the TJU dataset, the mean number of noise atoms per subject was 0.76 ± 0.95. We then repeated all primary analyses after excluding noise atoms. The resulting group-level maps were nearly identical to the original findings, with strong spatial concordance for both network correspondence (*r* = 0.93, spin-test–corrected *P* < 0.001) and *k*-hubness (*r* = 0.96, spin-test–corrected *P* < 0.001). Given the high reproducibility of results after noise exclusion, as well as the potential subjectivity introduced by manual noise classification, we report results based on the full, unfiltered set of SPARK atoms in the main analysis.

Second, to evaluate whether network correspondence findings generalized across alternative individualized network decomposition frameworks, we replicated the correspondence analysis using MELODIC independent component analysis (ICA) in FSL, one of the most widely used approaches for rs-fMRI network estimation. Consistent with prior work, the number of components was fixed at 30 for all subjects^25^. Focal epilepsy, as well as the TLE and EXE subgroups, showed a marked reduction in mean normativity and an increase in mean non-normativity when networks were derived using MELODIC-ICA (Supplementary Fig. 6). Correspondence patterns from MELODIC-ICA were highly similar to those obtained using SPARK in focal epilepsy (*r* = 0.50, spin-test–corrected *P* < 0.001; Supplementary Fig. 6), indicating that the correspondence findings are robust and not specific to a particular network decomposition method. Notably, because ICA enforces spatial independence among components, it does not support the estimation of overlapping networks and therefore precludes *k*-hubness analysis.

## DISCUSSION

Using an individualized network-estimation framework that permits overlapping intrinsic systems, together with multi-parcellation concordance and normative deviation modeling, we decomposed system-level functional topology into two complementary axes: system boundary fidelity captured by network correspondence (reduced normativity and increased non-normativity) and cross-system integration captured by *k*-hubness (redistribution of multi-network participation). This two-axis formulation provided a principled, patient-level description of network organization that is both interpretable and comparable across datasets. Importantly, these axes show dissociable clinical relevance: correspondence preferentially tracks neurocognitive deficits, whereas *k*-hubness more strongly reflects epilepsy clinical features.

Methodologically, our approach moves beyond node–edge connectivity summaries by defining measurements at the individualized system/network level. First, by estimating subject-specific intrinsic networks that explicitly permit overlap, we establish a substrate from which both (i) within-person system delineation can be derived from data-driven individualized networks rather than a priori templates, and (ii) cross-system participation can be quantified^17, 18, 19^. This allowed us to interpret *k*-hubness as multi-network participation that implied multi-functionality, rather than as a proxy for degree-based centrality within a fixed partition and singular functionality^8, 11, 22, 32^. Second, we leveraged multiple atlases spanning distinct parcellation principles and granularities to quantify individualized system integrity ^33, 34, 35^. This consensus-based strategy reduced dependence on any single atlas choice and enabled robust region-level quantification of correspondence. Third, *w*-score–based normative modeling anchored each patient’s topological profile to an age/sex- and covariate-adjusted healthy reference, yielding interpretable deviation maps that supported meaningful patient-level signatures and provided a principled basis for subsequent subtyping and staging ^8, 36, 37^.

Critically, multivariate sCCA indicated that while the network correspondence and cross-system integration axes do share variance, they remain uniquely dissociable. That is, correspondence-related deviations preferentially tracked neurocognitive variation, whereas *k*-hubness-related integrations more strongly aligned with epilepsy clinical features (e.g., lateralization, hippocampal sclerosis, and syndrome subtypes). These distinct linkages motivate our claim that these axes capture distinct topological phenotypes that implement different aspects of the epilepsy clinical condition. This unified multivariate result consolidates prior TLE-focused observations, where cognitive^25, 26^ and clinical^18, 19^ associations were examined in separate analyses, into a single coherent yet dissociable models within a focal epilepsy cohort that is then validated across a diverse non-focal sample.

In focal epilepsy, we observed a pervasive and directionally consistent disruption of network correspondence, marked by reduced normativity and increased non-normativity. This axis accords with prior TLE reports of diminished alignment to canonical system templates accompanied by increased atypical organization, underscoring correspondence as an interpretable index of system integrity ^25^. Its extension to EXE further suggested a shared mechanism of system-boundary degradation across focal epilepsies, with consistent expression across syndromic and lateralization strata (TLE and EXE; left- and right-lateralization). Notably, this disruption was not spatially random: within the consensus correspondence maps, the most prominent abnormalities localized to temporal, occipital, and frontal cortices, regions tightly linked to epileptic activity and its distributed network effects^12, 27, 38, 39, 40^. Biologically, this profile is most consistent with a progressive erosion of canonical system boundaries, whereby recurrent seizures and long-term plastic reorganization blur intrinsic network delineations and system identity, which may help explain why many studies observe widespread changes, variable across studies, in region-to-region coupling strength in focal epilepsy^8, 9, 10, 11, 20, 21^. Importantly, the spatial heterogeneity in correspondence we observed followed non-random and non-uniform disruption trajectories. Most notably, our SuStaIn analyses identified two reproducible progression patterns, distinguished by early involvement of pan-network/transmodal or predominantly unimodal systems, then converging on a common late-stage signature of widespread correspondence disruption. This convergence is consistent with connectome-gradient studies demonstrating atypical functional integration of transmodal and unimodal systems in TLE^10, 41, 42, 43^, and further suggested the presence of distinct “entry points” into network reorganization that ultimately funnel toward similar system-level outcomes as deterioration spreads. Importantly, these trajectories also imply partially distinct mechanisms with different temporal windows: early subtype-specific vulnerability of pan-network and transmodal organization may be most relevant for baseline cognitive risk, whereas later-stage correspondence disruption reflects a more generalized, cumulative functional reorganization that progresses largely independently of macroscopic gray matter loss.

Along the second axis, *k*-hubness delineated a syndrome-linked pattern of cross-system integration that was more spatially anchored to the epileptogenic network than the broadly shared correspondence endpoint. We found that focal epilepsy was characterized by reduced multi-network participation within temporal–limbic circuitry, most prominently in TLE, but also detectable in EXE. These transmodal, epileptogenic-network–colocalized regions^12, 27, 38, 39^ normally serve as cross-system “interfaces” by participating in multiple overlapping network, implying multi-functionality. Thus, decreased *k*-hubness suggested a loss of cross-system bridging capacity, with these nodes becoming more single-purpose or effectively confined to fewer systems. When interpreted alongside correspondence findings (i.e., altered canonical system boundaries in overlapping temporal–limbic territories), this interface loss is consistent with a more self-contained, intrinsically epileptogenic network organization that constrains information flow and limits flexible participation in the coordinated recruitment of distributed functional systems.^44, 45, 46^. In contrast, regions demonstrating increased *k*-hubness likely reflect heterogeneous biological processes rather than a unitary mechanism of reduced system integration. For instance, in association and sensory systems, including prefrontal and occipital cortices, *k*-hubness increases were observed consistent with compensatory redistribution of integrative load toward alternative pathways in response to focal system failure. This may serve as a way of preserving large-scale communication/integration when primary epileptogenic circuits lose connector capacity ^11, 18, 19, 21, 22^. In other contexts, the *k*-hubness increases we report may instead reflect epilepsy-driven network recruitment. For example, in extratemporal epilepsy, elevated *k*-hubness in primary motor regions may indicate enhanced integration of sensorimotor cortex into distributed epileptogenic networks, potentially facilitating seizure propagation or promoting maladaptive cross-system synchronization ^4, 10, 11, 47^. Together, these observations suggested that *k*-hubness increases can reflect either adaptive or maladaptive network reconfigurations, depending on anatomical context and syndrome-specific epileptogenic dynamics ^22^.

A central question is whether these system-level topological phenotypes simply index a generic “epilepsy effect” or whether they carry syndrome-specific information ^3, 5, 11, 27^. Cross-syndrome comparisons indicated both. That is, while GGE, SeLECTS, and absence epilepsy shared a partial overlap with focal epilepsy in correspondence and/or *k*-hubness abnormalities, their effects differed in direction, spatial distribution, and coherence. In GGE, correspondence deviations were comparatively diffuse and did not show the focal-typical temporo–fronto–occipital predominance observed in our focal epilepsy cohorts; whereas *k*-hubness did not exhibit a reproducible group-level shift. This divergence across the topological axes is consistent with large-scale evidence that focal and generalized epilepsies possess partially distinct macroscale organizational signatures^3, 5, 9, 27^. Notably, a similar minimal *k*-hubness signature was observed in absence epilepsy, whereas correspondence increases were preferentially expressed in thalamic and sensorimotor circuits. Consistent with models in which absence seizures arise from stereotyped thalamo–cortical spike–wave dynamics rather than multi-network connector involvement, these correspondence increases may reflect a more stereotyped expression of canonical thalamo–sensorimotor architecture^4, 11, 48, 49^. In contrast, SeLECTS showed increased thalamic correspondence alongside reduced cortical correspondence and *k*-hubness, producing a spatial profile distinct from adult focal epilepsy. This pattern is consistent with altered thalamo–cortical coupling in Rolandic circuits during development^50, 51^, and may reflect heightened vulnerability of maturing cortical systems that has been linked to increasingly recognized cognitive and learning difficulties in SeLECTS^52, 53, 54^. Importantly, our data clearly showed that these cross-syndrome differences are not well captured by a binary “focal versus generalized” dichotomy^3, 5, 9, 27^. Instead, our high-dimensional similarity analysis reframes the syndromes on a continuous spectrum: focal epilepsy occupies a separable region of feature space with larger deviations along both axes and a more distinctive topological pattern, while graded distances among syndromes (TLE ≈ EXE closer, GGE intermediate, SeLECTS and AE more distant) quantify a partial commonality alongside syndrome-specific heterogeneity in system-level organization ^2, 3, 22^.

Several limitations should be considered when interpreting these findings. First, our links to cognitive and clinical features are associational and cross-sectional. Confirming the network connectivity mechanisms implied by our system-level functional topologies, along with their neurocognitive and epilepsy-clinical feature linkages (as well as change over time), will require prospective outcome studies. In this regard, we note that our data remains silent with regard to the causal chain or direction of effects for the relationships we report between topology and the neurocognitive and clinical features. Second, while the present work established, in principle, the significance of both group-level shifts and patient-level variability in these topological biomarkers will require defined thresholds for abnormality, standardized patient-level reporting, and independent validation of their added value in the context of specific key clinical decisions and tasks (e.g., predicting surgical outcomes). Finally, although SuStaIn offered a principled framework for staging correspondence abnormalities, our staging inferences remain cross-sectional. Thus, longitudinal imaging will be necessary to confirm within-person progression, stage transitions, and their temporal relationship to clinical course and treatment.

In summary, our framework yielded two complementary, dissociable system-level topological phenotypes across the focal epilepsy spectrum. Correspondence disruptions appeared to be a broadly shared system-level endpoint that can be depicted at the patient level; whereas *k*-hubness reconfigurations are more closely aligned with epilepsy clinical features potentially implemented through lateralized and syndrome-dependent redistributions of cross-system integration. Together, these paired descriptions clarified how common network-level vulnerabilities and unique, specific topological reorganizations can coexist, providing interpretable, reproducible signatures for comparing individuals with regard to epilepsy network and brain system effects, as well as cognitive or treatment risks and comorbidities.

## MATERIALS AND METHODS

### Participants

A total of 1,208 presurgical epilepsy patients and 890 healthy participants (HPs) were included from two centers. The Thomas Jefferson University Comprehensive Epilepsy Center (TJU dataset) enrolled patients with focal epilepsy undergoing pre-surgical evaluation, including 305 focal epilepsy patients (age 15–73 years, 249 TLE, 56 EXE) and 224 HPs (age 18–65 years). The Jinling Hospital cohort (JLH dataset) aimed to recruit all consecutive visiting patients with common epilepsy syndromes, including 903 patients (age 6–56 years; 659 focal epilepsy, 108 GGE, 112 SeLECTS, 24 AE) and 666 HPs (age 4–70 years).

All patients underwent a comprehensive multidisciplinary evaluation including medical history, seizure semiology, continuous video-EEG telemetry, and clinical MRI, with PET performed when clinically indicated, to localize epileptogenic networks and establish syndrome classification^55, 56^. Key clinical variables of focal epilepsy patients were systematically documented, including epilepsy subtype (TLE, EXE), seizure lateralization (left, right, or uncertain), pathology category (unilateral hippocampal sclerosis, bilateral hippocampal sclerosis, other lesions, or normal MRI), and the occurrence of focal to bilateral tonic–clonic seizures (FBTCS) within the prior year.

In the TJU dataset, 276 patients completed a comprehensive neuropsychological assessment. Domains assessed included general verbal intellectual ability (Wechsler Adult Intelligence Scale [WAIS-III/IV]: IQ, Vocabulary, and Similarities), language (Controlled Oral Word Association Test: semantic and letter fluency; Boston Naming Test), executive function (WAIS-III/IV: Matrix Reasoning and Digit Span; Wisconsin Card Sorting Test: perseverative responses and categories completed; Trail Making Test Part B), mental efficiency and processing speed (Trail Making Test Part A; WAIS-III/IV: Digit Symbol Coding), verbal learning and memory (Wechsler Memory Scale-III/IV: Logical Memory I and II; California Verbal Learning Test: Total Learning and Long-Delay Free Recall), visuospatial ability (WAIS-III/IV: Block Design; Rey–Osterrieth Complex Figure Test, copy condition), and psychomotor functioning (Grooved Pegboard Test, dominant and non-dominant hands). Sample demographic and clinical characteristics are summarized in Supplementary Tables 1 (TJU) and 2 (JLH).

This study was approved by the Institutional Review Board for Research with Human Subjects at Thomas Jefferson University and at Jinling Hospital. All participants provided written informed consent.

### Imaging Methods

Thomas Jefferson hospital participants were scanned on a 3-T X-series Philips Achieva clinical MRI scanner (Amsterdam, the Netherlands) using an 8-channel head coil. A resting-state functional MRI scan lasting at least 5 minutes was acquired from all participants. During the resting-state condition, participants viewed a crosshair with no task requirements and were instructed to remain still throughout the scan and not fall asleep. After the scan, all participants reported that they were able to complete the scan as instructed. The fMRI data were collected with a single-shot echo-planar gradient echo imaging (EPI) sequence acquiring T2* signals (120-480 volumes; 34 axial slices acquired parallel to the anterior, posterior commissure line; TR = 2.5 s, TE = 35 ms; FOV = 256 mm, 128 × 128 data matrix voxels, flip angle = 90°, in-plane resolution = 2 mm × 2 mm, slice thickness = 4 mm). Each EPI imaging series started with three discarded scans to allow for signal stabilization. Prior to collection of the T2* images, T1-weighted images (180 sagittal slices) were collected using an MPRAGE sequence (256 × 256 isotropic 1mm voxels; slice thickness of 1 mm; TR = 640 ms; TE = 3.2 ms, FOV = 256 mm, flip angle = 8°) in positions identical to the functional scans to provide an anatomical reference.

Jinling hospital participants underwent functional and anatomical data acquisitions using a 3-T MRI scanner (SIEMENS Trio Tim, Siemens Healthcare). A resting-state functional MRI scan lasting at least 5 minutes was acquired from all participants. To minimize head movements, individuals were instructed to keep their eyes closed and remain awake during the scan, with a foam padding placed between their head and the coil. Functional MRI (fMRI) data were obtained using a single-shot echo-planar imaging sequence with the following parameters: TR = 2 s, TE = 35 ms, FOV = 240 mm, 64 × 64 data matrix voxels, flip angle of 90°, 30 transverse slices (slice thickness of 4 mm and interslice gap of 0.4 mm). T1-weighted images (176 sagittal slices) were collected using an MPRAGE sequence (256 × 256 isotropic 1-mm voxels; slice thickness of 1 mm; TR = 2300 ms; TE = 2.98 ms, FOV = 256 mm, flip angle = 90°) in positions identical to the functional scans to provide an anatomical reference.

### Data Preprocessing

Structural T1-weighted and functional data were first preprocessed with fMRIPrep v23.1.4^57^. Subsequent post-processing was performed using the eXtensible Connectivity Pipeline-DCAN (XCP-D) v0.7.3^58^, which applied standard nuisance regression and temporal filtering.

Specifically, nuisance regressors included the top five aCompCor components from white matter and CSF, six motion parameters and their temporal derivatives, and high-pass cosine regressors. The BOLD time series were denoised via linear regression, band-pass filtered (0.01–0.1 Hz), and spatially smoothed with a 6 mm FWHM Gaussian kernel. Finally, the denoised functional data were normalized to MNI152NLin2009cAsym space at 2 mm resolution.

After estimating head motion with fMRIPrep, we identified for each participant the continuous fMRI segment with the lowest motion that met a minimum length criterion (120 frames at TJU; 250 frames at JLH). Participants whose minimum-motion segment still showed mean head motion exceeding 0.4 (framewise displacement, FD; units as reported by fMRIPrep) were excluded from all following analyses.

### Individual-level Resting-state Functional Networks

We estimated individual resting-state functional networks using SParsity-based Analysis of Reliable *k*-hubness (SPARK) ^18, 19, 23, 24^, which integrates block-bootstrap resampling with a sparse GLM/dictionary-learning decomposition to recover overlapping resting state networks at the single-subject level. For each subject, the optimal number of networks was first determined on the original (non-bootstrapped) processed rsfMRI data by searching across 18 to 40 components in increments of two. The subject’s rsfMRI time series was then resampled B = 200 times using temporal blocks of 20–60 s, a procedure that preserves low-frequency signal structure and autocorrelation. For each bootstrap realization, SPARK learns a data-driven temporal dictionary together with a sparse spatial coefficient map, such that each voxel’s time series is represented as a sparse linear combination of a small number of network “atoms.” These atoms correspond to functional networks that are explicitly allowed to overlap in space. The resulting ensemble of spatial maps across bootstraps is clustered to yield consensus networks, after which reproducibility filtering is applied to retain only the stable components. The retained set constitutes each subject’s reliable resting-state networks. Notably, SPARK has been shown to exhibit high single-subject test–retest reliability^18^, providing a reliable substrate for topological and *k*-hubness analyses. For computational efficiency, SPARK was performed on rsfMRI downsampled to 4-mm isotropic resolution; final subject-specific networks were then resampled to 2-mm MNI space for subsequent correspondence. All spatial atoms identified by SPARK were visually inspected by a physician with training in radiology and neuroscience, following established criteria for identifying noise components (e.g., atom located along brain boundaries or outside the brain, as well as vascular- and motion-related artifacts) ^31^. To account for potential subjectivity, analyses were repeated in parallel using both the full set of atoms and the subset after noise exclusion.

In addition to SPARK, we derived individual resting-state networks using MELODIC-ICA in FSL, with the model order fixed at 30 components as in previous work ^25^. These 30 spatially independent networks per subject were used to validate the networks obtained with SPARK.

### Network Correspondence Profiles

We assessed the normativity of individualized functional network topology by quantifying spatial correspondence between each subject’s individualized network set (K) and canonical intrinsic connectivity networks (ICNs; M) defined by multiple widely used cortical and subcortical atlases^25^. For each subject and atlas, we computed a K × M overlap matrix and summarized it into subject-level indices of topological alignment and deviation.

#### Cortical topology (surface-based correspondence)

Pairwise overlap between individualized and canonical networks was computed using Network Correspondence Toolbox ^34^ which projects subject-specific networks from MNI space to the fsaverage6 cortical surface and calculates Dice similarity coefficients for all K × M network pairs, yielding a subject-level similarity matrix. Statistical significance of these correspondences was assessed using 1,000 spin-based spatial permutations, which test whether observed overlaps exceed those expected under spatially rotated null distributions^59^. To ensure robustness, we repeated analyses across multiple canonical atlases that vary in network number and parcellation strategy, including AS200Y17 (Alex Schaefer200-parcels ^28^ with Yeo 17 networks^60^), EG17 (Gordon 17 networks ^15^), HCP-ICA (25-component ICA maps from the Human Connectome Project ^61^), MG360J12 (Glasser 360 parcels ^62^ mapped to Ji 12 Cole–Anticevic networks ^63^), TY7 (Thomas Yeo 7 networks^60^), and UKB-ICA (25-component ICA maps from the UK Biobank ^64^).

#### Subcortical topology (volumetric correspondence)

For subcortical topology, we computed Dice overlap in MNI space between individualized networks and subcortical structures delineated by the FreeSurfer ASEG segmentation ^65^, encompassing seven bilateral regions. As spin permutation testing is not applicable to volumetric data, statistical inference was not performed for subcortical correspondences.

#### Atlas-specific normativity metrics

For each subject and atlas, let D ∈ ℝ^K×M^ denote the Dice similarity matrix where D_k,m_is the Dice overlap between individualized network k and canonical ICN m. From D, we derived three subject-level metrics:

**1. Canonical Network Representation (CNR)**: For each canonical ICN m (i.e., each column of the K × M matrix), we selected the maximum Dice overlap across the K individualized networks (column-wise maximum).

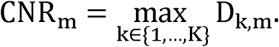

This yields an M-length vector per subject and atlas, capturing how strongly each canonical ICN is expressed in that individual (via its closest individualized counterpart).

**2. Normativity**: We defined overall (global) normativity as the mean CNR across canonical ICNs in that atlas:

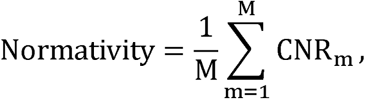

capturing the overall fidelity with which an individual’s functional network topology conforms to canonical network configurations.

**3. Non-normativity:** For each individualized network k, we computed the fraction not explained by any canonical ICN (one minus the row-wise maximum), then averaged across individualized networks:

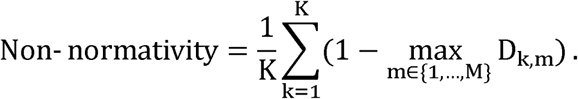

This metric indexes topological deviation or idiosyncrasy relative to canonical network architecture.

#### Consensus correspondence map across canonical atlases

To obtain a consensus estimate of how consistently each subject expresses canonical functional systems across atlases (rather than relative to any single atlas), we constructed a cross-atlas consensus representation using cortical atlases. For each subject and atlas, we first started from the atlas-specific CNR vector (best-match Dice values for each canonical network), where all atlas-specific representations were mapped to a common cortical reference by expressing correspondence in AS200 parcels defined on the Conte69/fs_LR_32k surface. This consensus-mapping step operates on atlas-level CNR vectors (network-wise scalars) and uses canonical atlas geometry only to distribute these scalars into parcel space.

For label-based atlases (AS200Y17, EG17, MG360J12, TY7), canonical network labels were available at the vertex level in fs_LR_32k space. For each atlas, we constructed a network-to-parcel weight matrix W ∈ ℝ^M×200^ where each entry W_m,p_ equals the number of vertices jointly belonging to canonical network m and parcel p, normalized by the total number of vertices in parcel p. Given a subject’s CNR vector c ∈ ℝ^M^x, the atlas-specific parcel-wise correspondence profile was computed as:

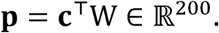

When a parcel overlapped multiple canonical networks, its correspondence value therefore reflected a weighted average of the corresponding CNR values, with weights proportional to the fraction of parcel vertices assigned to each network.

For ICA-based atlases (HCPICA and UKBICA), canonical “networks” are spatial components that may partially overlap. We defined each ICA component’s vertex-wise support mask on the fs_LR_32k surface as vertices with component weight > 0. A Schaefer-200 parcel was considered covered by a component if any of its vertices were included in that component’s >0 mask. For each subject and parcel, the atlas-specific parcel value was computed as the mean CNR across all ICA components covering that parcel. Parcels not covered by any component were assigned zero and were excluded from across-atlas averaging for that parcel.

Within this shared parcel space, we generated a subject-specific **consensus correspondence map** by averaging parcel-wise values across atlases for each parcel, using only atlases that provided coverage for that parcel. This consensus profile reflects the stability of canonical network expression across heterogeneous atlas definitions: parcels with high consensus values indicate regions where an individual’s networks showed consistently strong canonical alignment across parcellation- and ICA-based definitions, whereas low values indicate regions where correspondence is weak or atlas-dependent, highlighting potentially individual-specific organization or boundary ambiguity. Finally, to obtain a network-level summary, the parcel-wise consensus profile was averaged within Yeo-17 networks by mapping Schaefer-200 parcels to the Yeo-17 partition and computing the mean across parcels within each network.

### *k*-hubness (Multi-Network Participation) profiles

From the SPARK-derived overlapping networks, we quantified voxel-wise *k*-hubness as the number of consensus networks in which a voxel exhibited non-zero loading (i.e., participation). A voxel with *k* = 1 participates in only a single network and is therefore considered a non-hub in this framework, whereas a voxel with *k* ≥ 2 participates in multiple networks and is indicative of a connector hub. Connector hubs are regions that link distinct functional systems through multi-network participation, thereby supporting communication and integration across the whole brain. In this way, *k*-hubness indexes the extent to which a voxel serves as an interface across functional systems rather than being confined to a single network^18, 19^.

#### Network/region *k*-hubness

We summarized voxel-wise *k*-hubness within predefined cortical and subcortical networks or regions by averaging across voxels. All summaries were restricted to gray-matter masks. This index captures the degree to which a given network or region participates in multiple overlapping systems. Cortical partitions followed the AS200Y17 parcellation^28, 60^; to accommodate potential lateralization, the 17 canonical networks were split into left and right homologues, yielding 34 cortical networks. Subcortical regions were defined using FreeSurfer ASEG segmentation^65^ and analyzed separately for seven regions per hemisphere.

### Normative Model (*w*-score)

We used PCNtoolkit ^37^ to convert all network/region metrics into individualized deviation scores (*w*-scores). Within each cohort, we trained the model exclusively on that cohort’s healthy participants to learn the expected value of each metric as a function of sex, age, head motion, and the number of SPARK atoms; the resulting cohort-specific models were then applied to all individuals within the same cohort to quantify deviation from the normative pattern. For each metric, we fit a Bayesian linear regression normative model with heteroskedastic noise and a B-spline basis to capture smooth, non-linear covariate effects. In practice, we used a **cubic B-spline** with **four knots** distributed across the covariate range, providing enough flexibility to model gentle curvature while avoiding overfitting. The fitted model returned, for every participant, an expected value and a residual; we expressed the residual as a *w*-score by scaling it to the residual variability observed in the healthy group. These *w*-scores are reported in units of typical healthy variation, retain the sign of deviation, and serve as a harmonized effect size across metrics and cohorts.

### Statistical analysis

All analyses were performed on covariate-adjusted *w*-scores, with statistical inference testing deviations from the normative reference (w = 0). Different multiple-comparison correction strategies were applied depending on the dimensionality of the metric. For global, whole-brain summary indices (e.g., Normativity and Non-normativity), group-level deviations were assessed using two-sided one-sample t tests against zero, and between-group multiple comparisons were controlled using the Benjamini–Hochberg false discovery rate (BH-FDR) procedure.

For high-dimensional network- or parcel-resolved measures, including consensus correspondence maps and *k*-hubness profiles, group-level deviations were assessed using permutation based two-sided one-sample t tests (10,000 permutations) against zero ^66, 67^, with max-T correction applied across networks or parcels. Permutation-based inference was implemented using PERMUTOOLS^68^ (https://github.com/mickcrosse/PERMUTOOLS). This approach controls the family-wise error rate by comparing observed test statistics to the maximum statistic obtained under the null distribution across all regions.

### Subtype and stage inference (SuStaIn)

To model latent disease heterogeneity and infer putative progression patterns from cross-sectional neuroimaging abnormalities, we applied the Subtype and Stage Inference (SuStaIn) framework ^12, 14, 69^ to individualized deviation measures derived from normative modeling. SuStaIn conceptualizes disease evolution as a mixture of subtype-specific trajectories, in which biomarkers transition monotonically from normality to increasing abnormality across a sequence of discrete stages. This modeling assumption makes SuStaIn particularly well suited for features that plausibly reflect cumulative pathological burden. Based on this principle, we constructed separate SuStaIn models using two complementary feature sets: GMV abnormalities and correspondence-based network measures. Both feature families demonstrated approximately monotonic changes with disease burden at the group level, consistent with the core assumptions of the SuStaIn framework. Note, *k*-hubness-related metrics were not included in SuStaIn modeling as *k*-hubness reflects complex, non-monotonic reconfigurations of network topology that may capture either compensatory (maladaptive) or adaptive processes rather than unidirectional pathological progression.

For network correspondence based SuStaIn analysis, features were derived from two global correspondence indices (normativity and non-normativity), network-level correspondence measures for the 17 canonical cortical networks defined by the AS200Y17 atlas, and CNRs for seven pairs of bilateral subcortical regions, yielding a total of 26 correspondence features.

In parallel, for the GMV-based SuStaIn analysis, regional features were derived from bilateral cortical networks defined by the AS200Y17 atlas (34 networks), together with 14 subcortical regions. All GMV measures were first adjusted for relevant covariates (sex, age, total intracranial volume) using normative modeling to obtain subject-specific deviation scores. To limit model complexity and reduce redundancy among highly correlated regions, we performed anatomically and functionally informed feature aggregation prior to SuStaIn fitting. Specifically, homologous or closely related cortical networks and subcortical structures were combined into composite features, yielding a final GMV feature set of 28 regions. This dimensionality reduction strategy improved model stability and interpretability while preserving sensitivity to spatially distinct patterns of atrophy. The full mapping between original regions and aggregated features is provided in the Supplementary Table 4.

To ensure comparability between correspondence and GMV analyses, we restricted this SuStaIn analysis to participants with high-quality T1-weighted structural scans, excluding seven individuals with motion-related artifacts. All features were adjusted for relevant covariates using normative modeling, resulting in subject-specific *w*-scores that quantify deviations from the healthy reference distribution. To ensure consistency with the SuStaIn assumption of monotonic progression, features were sign-aligned based on their sample-level mean direction such that larger values corresponded to increasing abnormality with disease progression. SuStaIn models were fitted using a *z*-score–based event formulation, in which each feature transitions through increasing abnormality levels across stages. Fixed *z*-score waypoints of 1, 2, and 3 were used. Models with one to four candidate subtypes were fitted using 25 random start points and 10,000 Markov chain Monte Carlo iterations. The optimal number of subtypes was determined using five-fold cross-validation based on the cross-validation information criterion (CVIC) and out-of-sample log-likelihood, after which the selected model was refit using the full dataset. For each individual, the final model yielded posterior probabilities over subtype-by-stage combinations, from which maximum-likelihood subtype membership and disease stage assignments were derived.

### Sparse Canonical Correlation Analysis(sCCA)

We applied sCCA (PMA v.1.2.4 package in R) ^70^ to relate a combined network topology feature set (pooled network-correspondence and *k*-hubness) to a combined clinical–behavioral set comprising epilepsy clinical features (seizure type, lateralization, pathology, FBTCS) and cognitive measures (language–memory, executive functions, motor skills, and related domains). The number of canonical dimensions was constrained a priori and capped at four. Sparsity penalties for the imaging and clinical sides were selected in a data-driven manner using CCA.permute function with 200 permutations, and the selected penalty ratios were then used to fit the final sCCA model. To assess statistical significance, a permutation p-value was obtained with MultiCCA.permute function using 1000 permutations. For each retained dimension, we report the canonical correlation and summarize feature contributions on both sides via the non-zero loadings and their associated weights.

## Acknowledgments

The authors thank all the healthy individuals and patients with epilepsy who participated in this study. This work utilized computational resources provided by the Advanced Cyberinfrastructure Coordination Ecosystem: Services & Support (ACCESS), supported by the National Science Foundation (MED050005).

## Data availability

Anonymized patient level imaging and clinical metrics are available on the Zenodo at https://doi.org/10.5281/zenodo.18808544.

## Code availability

All analysis code used in this study is publicly available at https://github.com/qiruizhangqirui/epilepsy-network-topology. Functional MRI preprocessing was performed using fMRIPrep (v23.1.4; https://fmriprep.org/en/stable/) and XCP-D (v0.7.3; https://xcp-d.readthedocs.io). Individual resting-state functional networks were estimated using SPARK (https://github.com/multifunkim/spark-matlab) or MELODIC-ICA implemented in FSL (https://fsl.fmrib.ox.ac.uk/fsl/docs/). Pairwise overlap between individualized and canonical networks was computed using the Network Correspondence Toolbox (https://github.com/rubykong/cbig_network_correspondence). Normative modeling and *w*-score computation were performed using PCNtoolkit (https://github.com/amarquand/PCNtoolkit).

Permutation-based statistical testing was conducted using PERMUTOOLS (https://github.com/mickcrosse/PERMUTOOLS). Subtype and stage inference analyses were performed using pySuStaIn (https://github.com/ucl-pond/pySuStaIn). Sparse canonical correlation analysis (sCCA) was implemented using the PMA package (v1.2.4; https://github.com/cran/PMA). Brain surface visualizations were generated using the ENIGMA Toolbox (https://enigma-toolbox.readthedocs.io/).

