## Supplementary Fig. 1 for "Mapping Individualized Dual-Axis Network Topology in Focal Epilepsy: Divergent Alterations in System Integrity, Integration, and Clinical Correlates"

Supplementary Table 1. Sample demographic and clinical characteristics of TJU dataset

| TJU dataset | **Focal epilepsy**  **(*n* = 305)** | **Healthy participants**  **(*n* = 224)** |
| --- | --- | --- |
| Age  (range) | 39.23 ± 13.25  (15-73) | 32.80 ± 10.36  (18-65) |
| Sex (Male/Female) | 153/152 | 113/111 |
| Age at epilepsy onset (years) | 23.74 ± 14.78 |  |
| Duration of epilepsy (years) | 15.49 ± 13.99 |  |
| Epilepsy type (TLE/EXE) | 249/56 |  |
| Seizure type (with/without FBTCS) | 187/118 |  |
| Seizure lateralization  (Left/Right/Uncertain) | 159/116/30 |  |
| Pathology (NB/UHS/BHS/Lesion) | 86/104/25/90 |  |
| fMRI head motion (FD) | 0.15 ± 0.07 | 0.13 ± 0.06 |

Continuous variables are presented as mean ± SD.

Pathology was determined based on neuroimaging: NB = normal brain; UHS = unilateral hippocampal sclerosis; BHS = bilateral hippocampal sclerosis; FBTCS = focal to bilateral tonic-clonic seizures.

Supplementary Table 2. Sample demographic and clinical characteristics of JLH dataset

| JLH dataset | **Focal epilepsy**  **(*n* = 659)** | **GGE**  **(*n* = 108)** | **SeLECTS**  **(*n* = 112)** | **Absence**  **epilepsy**  **(*n* = 24)** | **Healthy participants**  **(*n* = 666)** |
| --- | --- | --- | --- | --- | --- |
| Age  (range) | 26.45 ± 9.73  (6-56) | 25.35 ± 8.40  (14-52) | 9.31 ± 2.24  (5-15) | 10.33 ± 3.96  (6-21) | 33.30 ± 17.78  (4-70) |
| Sex (Male/Female) | 357/302 | 75/33 | 55/57 | 10/14 | 296/370 |
| Epilepsy type (TLE/EXE) | 405/254 |  |  |  |  |
| Seizure type  (with/without FBTCS) | 505/154 |  |  |  |  |
| Seizure lateralization  (Left/Right/Uncertain) | 202/185/272 |  |  |  |  |
| Pathology (NB/UHS/BHS/Lesion) | 240/177/32/210 |  |  |  |  |
| fMRI head motion (FD) | 0.19 ± 0.07 | 0.18 ± 0.08 | 0.18 ± 0.07 | 0.16 ± 0.05 | 0.19 ± 0.08 |

Continuous variables are presented as mean ± SD. GGE = genetic generalized epilepsy; SeLECTS = self-limited epilepsy with centrotemporal spikes.

Pathology was determined based on neuroimaging: NB = normal brain; UHS = unilateral hippocampal sclerosis; BHS = bilateral hippocampal sclerosis; FBTCS = focal to bilateral tonic-clonic seizures.

Supplementary Table 3. Aggregated features for GMV-based SuStaIn analysis.

| Original regions | Aggregated regions | Original regions | Aggregated regions |
| --- | --- | --- | --- |
| Left_TempPar | Left_TempPar | Right_TempPar | Right_TempPar |
| Left_DefaultC | Left_DefaultC | Right_DefaultC | Right_DefaultC |
| Left_DefaultB | Left_DefaultB | Right_DefaultB | Right_DefaultB |
| Left_DefaultA | Left_DefaultA | Right_DefaultA | Right_DefaultA |
| Left_ControlC |  | Right_ControlC |  |
| Left_ControlB | Left_Control | Right_ControlB | Right_Control |
| Left_ControlA |  | Right_ControlA |  |
| Left_LimbicA | Left_Limbic | Right_LimbicA | Right_Limbic |
| Left_LimbicB |  | Right_LimbicB |  |
| Left_Sal_VenAttnB | Left_Sal_VenAttn | Right_Sal_VenAttnB | Right_Sal_VenAttn |
| Left_Sal_VenAttnA |  | Right_Sal_VenAttnA |  |
| Left_DorsAttnB | Left_DorsAttn | Right_DorsAttnB | Right_DorsAttn |
| Left_DorsAttnA |  | Right_DorsAttnA |  |
| Left_SomatomotorB | Left_Somatomotor | Right_SomatomotorB | Right_Somatomotor |
| Left_SomatomotorA |  | Right_SomatomotorA |  |
| Left_VisualB | Left_Visual | Right_VisualB | Right_Visual |
| Left_VisualA |  | Right_VisualA |  |
| Left_Caudate | Left_Striatum | Right_Caudate | Right_Striatum |
| Left_Putamen |  | Right_Putamen |  |
| Left_Hippocampus | Left_Hippocampus | Right_Hippocampus | Right_Hippocampus |
| Left_Pallidum | Left_Pallidum | Right_Pallidum | Right_Pallidum |
| Left_Thalamus | Left_Thalamus | Right_Thalamus | Right_Thalamus |

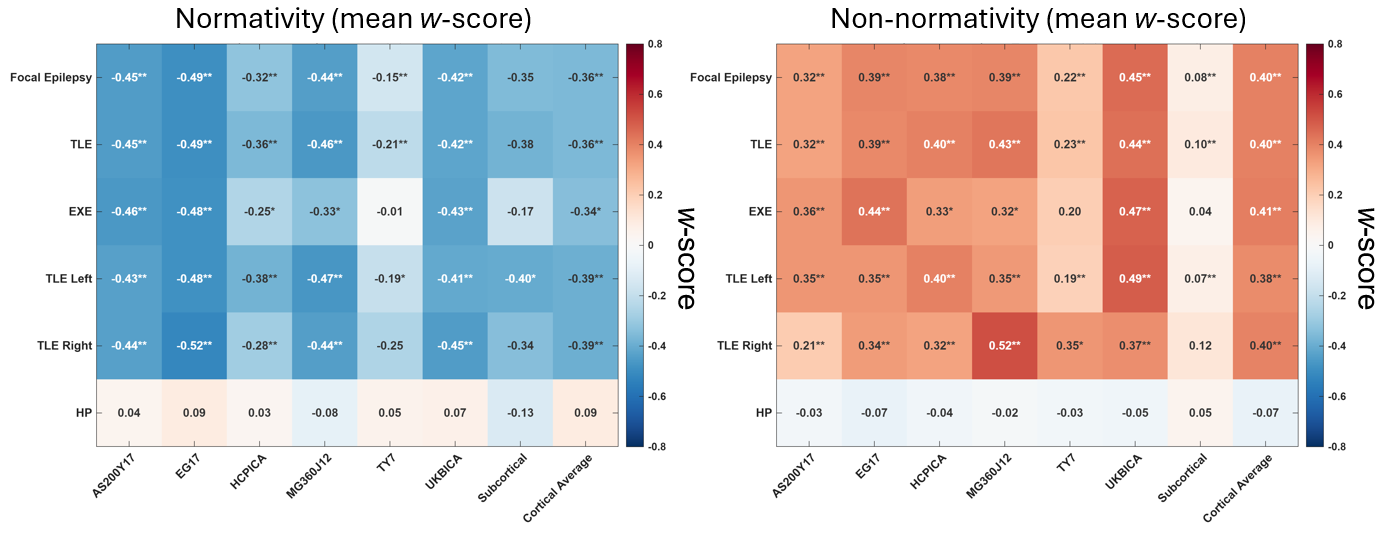

**Supplementary Figure 1. Atlas-specific patterns of normativity and non-normativity.**
Heatmaps display the *w*-scores and statistical significance of normativity (left) and non-normativity (right) for healthy participants (HP) and each focal epilepsy subtype across all canonical network atlases. Across atlases, the direction and relative magnitude of effects are consistent with the aggregated results shown in Figure 2, confirming reduced normativity and increased non-normativity in focal epilepsy. Notably, subcortical non-normativity alterations are minimal (normativity is also reduced, though to a lesser extent), suggesting that non-normative network organization is predominantly expressed cortically rather than subcortically.

**P* < 0.05, ***P* < 0.01, FDR-corrected.

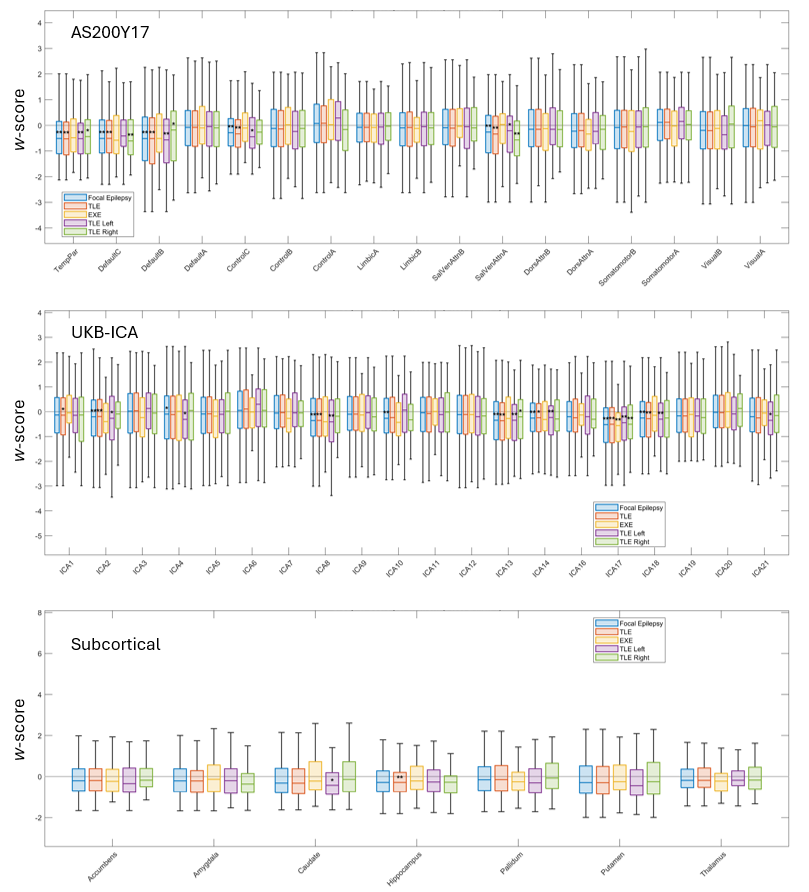

**Supplementary Figure 2. Atlas-specific canonical network representation (CNR).**
Atlas-specific patterns of CNR are shown for three representative atlases: AS200Y17, a functionally defined hard-parcellation atlas; UKB-ICA, an ICA-derived soft-parcellation atlas; and FreeSurfer ASEG, a subcortical atlas. The mean value for HPs is centered at zero on the y-axis. Across these atlases, CNR results demonstrate network correspondence disruptions consistent with those observed in Figure 2. Statistical significance is assessed within each group across networks using permutation-based max-T correction. **P* < 0.05, ***P* < 0.01.

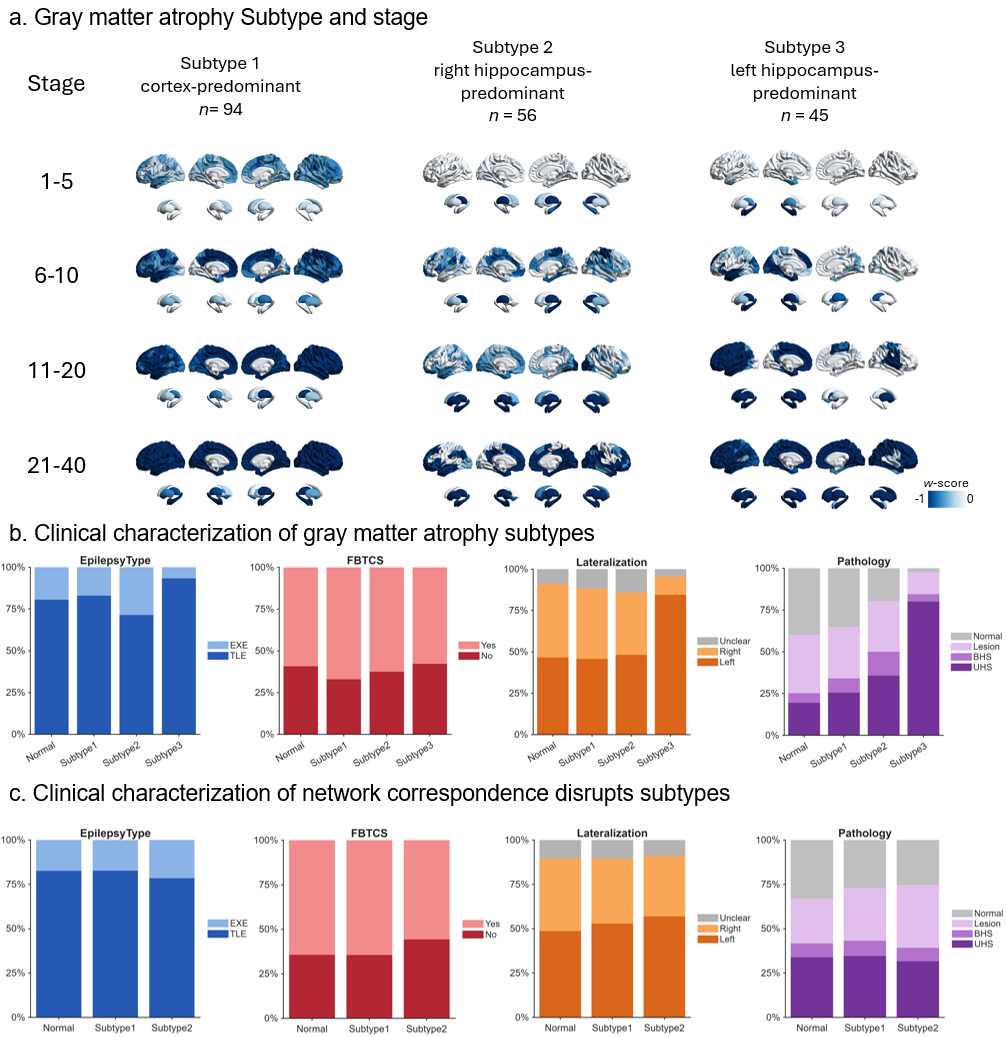

**Supplementary Figure 3. Subtypes and staging of gray matter atrophy in focal epilepsy. (a)** Mean regional *w*-scores of gray matter atrophy across inferred stages for the three SuStaIn-derived subtypes: cortex-predominant, right hippocampus–predominant, and left hippocampus–predominant. **(b)** Clinical characterization of gray matter atrophy subtypes. Epilepsy type differs across subtypes (χ² = 8.15, *P* = 0.042), with a higher proportion of TLE in the left hippocampus–predominant subtype. Differences in focal-to-bilateral tonic–clonic seizures (FBTCS) were not significant (χ² = 1.69, *P* = 0.063). Seizure lateralization differs markedly across subtypes (χ² = 23.68, *P* < 0.0001), with a higher proportion of left-sided seizures in the left hippocampus–predominant subtype. Pathology also differs significantly (χ² = 64.37, *P* < 0.001), with both right and left hippocampus–predominant subtypes showing higher proportions of unilateral and bilateral hippocampal sclerosis. **(c)** Clinical characterization of network correspondence disruption subtypes. Correspondence subtypes showed no significant differences across clinical variables, including epilepsy type (χ² = 0.67, *P* = 0.715), FBTCS (χ² = 1.86, *P* = 0.394), seizure lateralization (χ² = 1.38, *P* = 0.847), or underlying pathology (χ² = 3.03, *P* = 0.808). BHS: bilateral hippocampal sclerosis; UHS: unilateral hippocampal sclerosis.

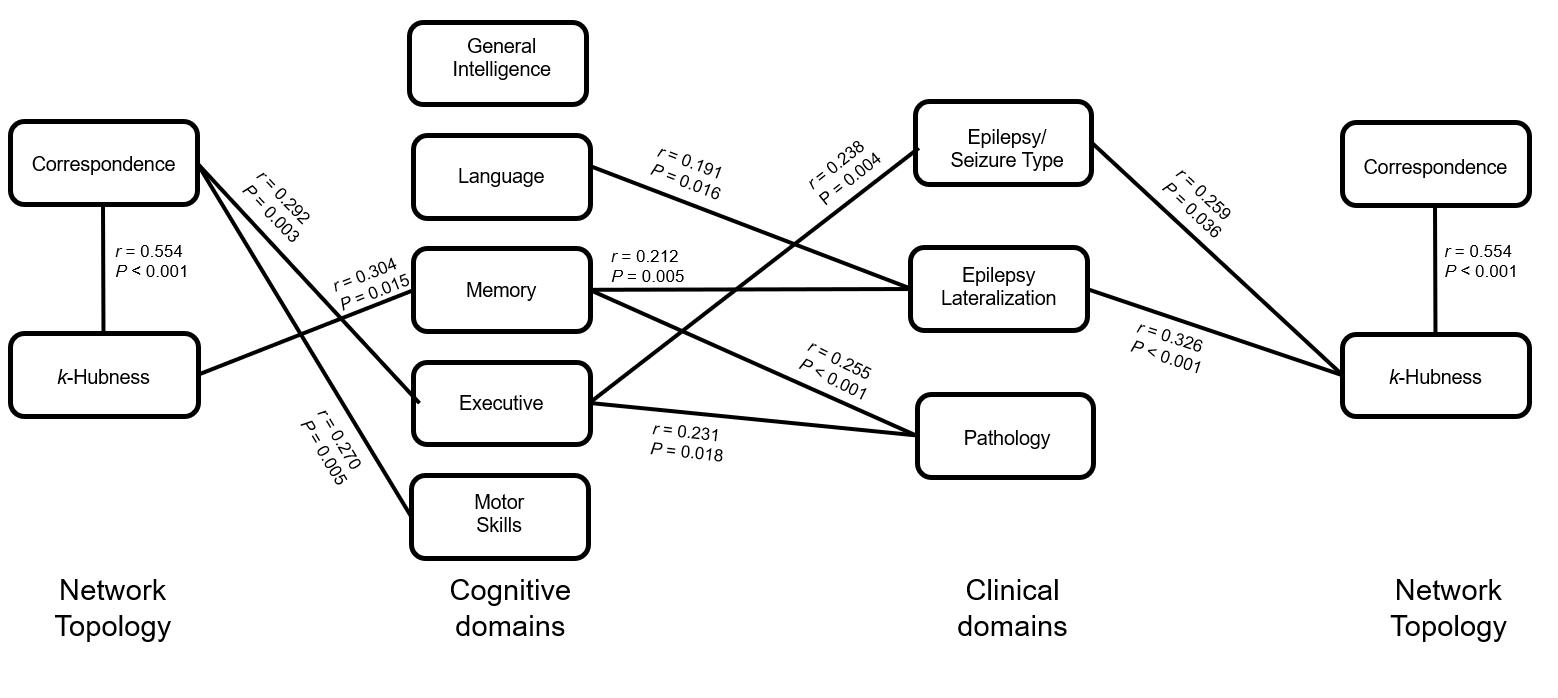

**Supplementary Figure 4. Fine-grained subdomain-level sparse canonical correlation analysis of network topology and clinical–cognitive phenotypes.** Subdomain-level sCCA is used to resolve domain-specific associations between network topology and clinical–cognitive phenotypes. r values denote the canonical correlation for Variate 1 from each pairwise sCCA. Lines indicate significant associations identified by permutation testing (*P* < 0.05). This analysis is exploratory and intended to contextualize and refine the main multivariate findings; therefore, no multiple-comparison correction across domain pairs is applied. Across cross-domain associations, correspondence-profile features (System Integrity) are most strongly associated with executive (*r* = 0.292, *P* = 0.003) and motor (*r* = 0.270, *P* = 0.005) performance. In contrast, *k*-hubness features (System Integration) align more strongly with epilepsy subtype/seizure type (latent construct including TLE vs. EXE and focal-to-bilateral tonic–clonic seizures [FBTCS], *r* = 0.259, *P* = 0.036), seizure lateralization (latent construct including left, right, and uncertain, *r* = 0.326, *P* < 0.001), and memory performance (*r* = 0.304, *P* = 0.015). Within-domain associations are also evident. Correspondence profiles are partially explained by *k*-hubness features (*r* = 0.554, *P* < 0.001), indicating shared variance across the two topology axes. Language (*r* = 0.191, *P* = 0.016) and memory (*r* = 0.212, *P* = 0.005) covary with lateralization; executive performance tracks seizure type (*r* = 0.238, *P* = 0.004); and network–phenotype associations for memory (*r* = 0.255, *P* < 0.001) and executive function (*r* = 0.231, *P* = 0.018) are modulated by underlying pathology.

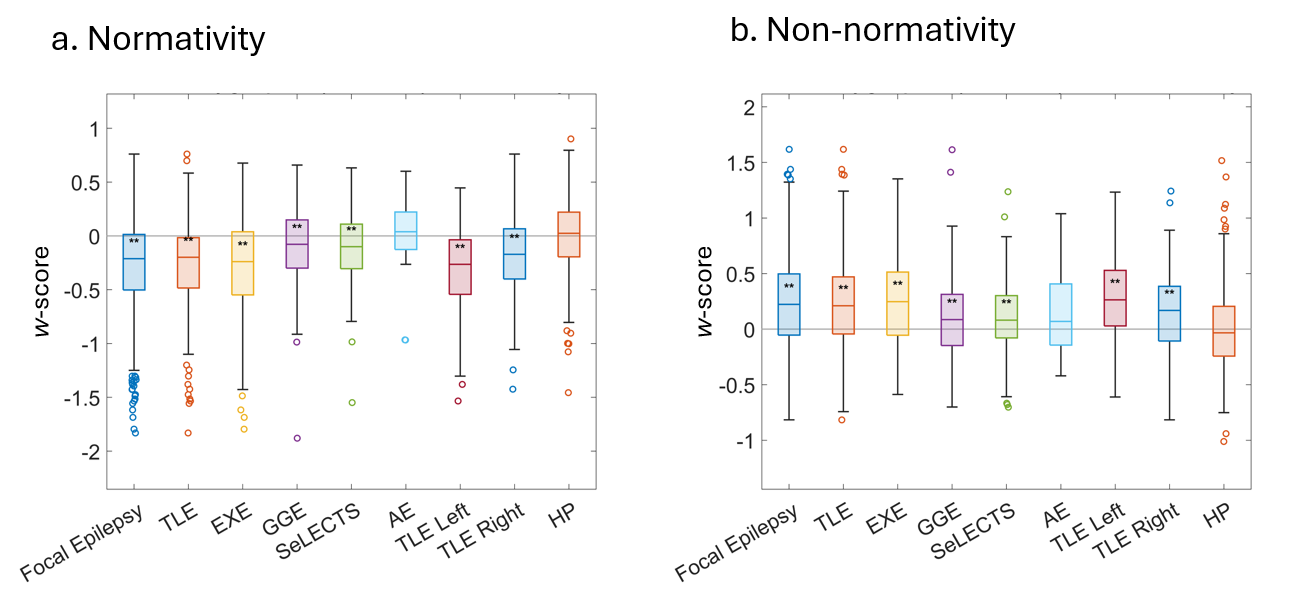

**Supplementary Figure 5. Normativity and Non-normativity in the JLH cohort.**
Replication of normativity and non-normativity in an independent cohort (JLH), extended to additional common epilepsy syndromes. (a) Across canonical atlases, individuals with focal epilepsy, TLE, EXE, GGE, SeLECTS, absence epilepsy, as well as left- and right-sided TLE exhibit a marked reduction in mean normativity. (b) Conversely, the same groups show a significant increase in mean non-normativity across atlases.

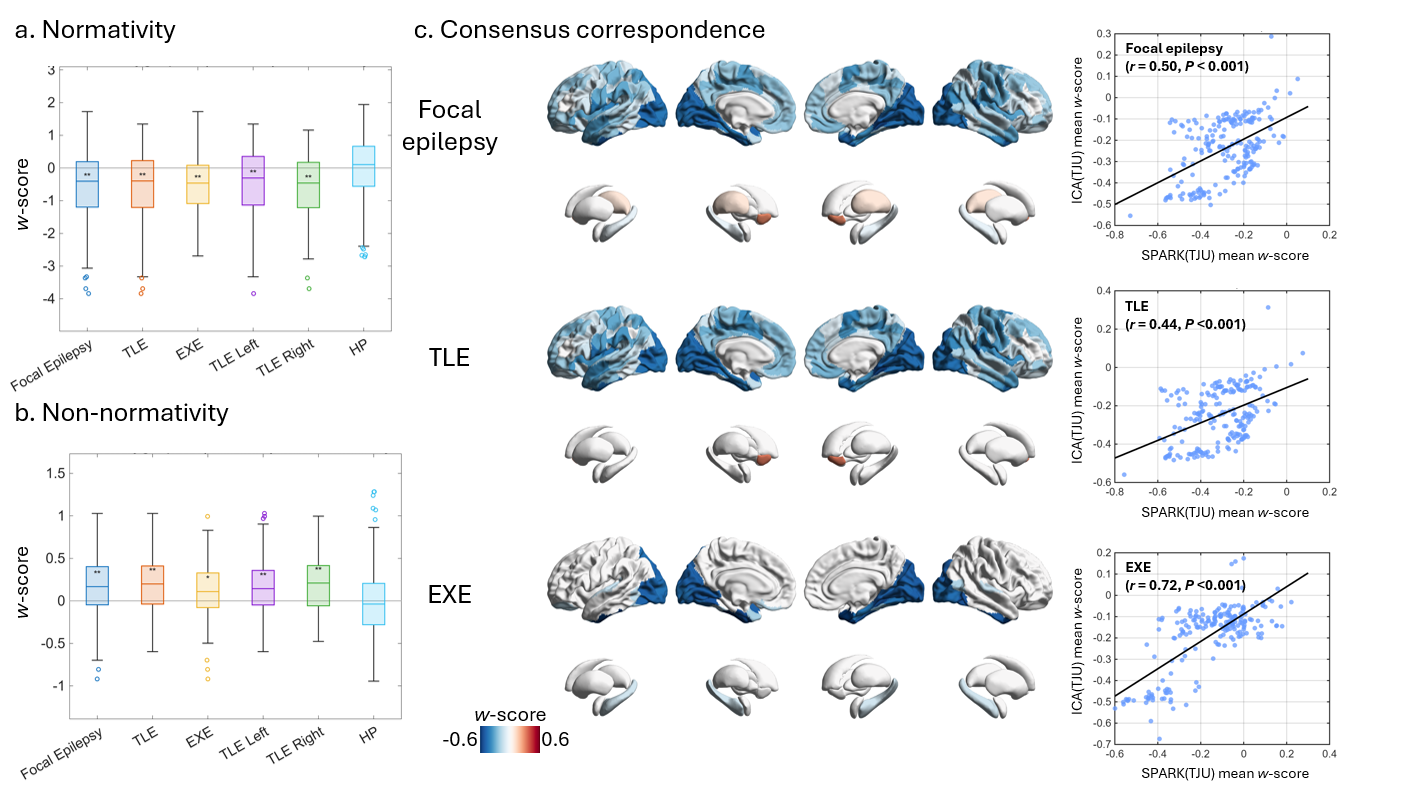

**Supplementary Figure 6. Replication of network correspondence findings using ICA-based individualized networks.**
To assess the generalizability of network correspondence results across alternative individualized network decomposition methods, correspondence analyses are replicated using MELODIC independent component analysis (ICA) implemented in FSL, with the number of components fixed at 30 for all subjects. **(a)** Group differences in mean normativity and **(b)** mean non-normativity *w*-scores derived from ICA-based individualized networks(**P* < 0.05, ***P* < 0.01, FDR corrected). **(c)** Spatial consensus correspondence maps (*w*-scores) aggregated across atlases, highlighting cortical regions with significantly reduced correspondence in focal epilepsy and in each subgroup (*P* < 0.05).

For each group, the accompanying scatter plots show the spatial correlation between correspondence maps derived from ICA-based and SPARK-based network decompositions (Pearson correlation; spin-test-corrected), demonstrating high spatial concordance between methods.
